# CROP2, a Retriever-PROPPIN Complex Mediating Protein Export from Endosomes to the Plasma Membrane

**DOI:** 10.1101/2025.10.08.681146

**Authors:** Maria Giovanna De Leo, Andreas Mayer

## Abstract

Endosomes generate tubulo-vesicular carriers to redistribute proteins between plasma membrane, Golgi, and lysosomes. These transport routes employ distinct combinations of sorting nexins with complexes such as Retromer or Retriever. We now show that, while Retromer associates with the PROPPIN WIPI1 to form the previously described CROP complex, Retriever associates with WIPI2, forming CROP2. WIPI2 integrates into Retriever-dependent coat complexes, since it interacts both with the Commander subunit CCDC93 and its cognate sorting nexin SNX17. CROP and CROP2 are exclusive in their physical associations and pathway selective. Whereas CROP2 is required for endosomal exit of β1-Integrin, it does not affect CROP-dependent cargos, such as EGFR or GLUT1. Vice versa, CROP is not required for β1-Integrin trafficking. WIPI1 and WIPI2 rely on similar molecular features. Their activity depends on the same FSSS motif to integrate into Retromer and Retriever complexes, respectively, and on an amphipathic membrane-inserting α-helix, which conveys membrane fission activity to PROPPINs. This suggests that Retromer and Retriever coats integrate distinct PROPPIN isoforms to promote fission of the respective endosomal carriers formed by them.

## Introduction

Endosomal membrane traffic regulates the abundance and activity of many receptors and transporters at the plasma membrane, which renders it critical for the development and homeostasis of cells and tissues (McNally and Cullen, 2018). In line with this, perturbations in endosomal trafficking contribute to a wide range of diseases, including cancer and some of the most common neurological disorders (Carosi et al., 2023; Mellman and Yarden, 2013; Schreij et al., 2016). Many plasma membrane proteins cycle between the cell envelope, endosomes, and the Golgi. They can move on to lysosomes for degradation unless they are retrieved before endosomes fuse with lysosomes (Cullen and Steinberg, 2018). Endosomes thus serve as sorting stations at an intersection between several compartments. To collect cargo proteins, endosomes employ a variety of coats that form tubulo-vesicular carriers. Many of them are based on sorting nexins, which scaffold the membrane into narrow tubules. Sorting nexins can act on their own, such as in the ESCPE-1 complex (Kvainickas et al., 2017; Lopez-Robles et al., 2023; Simonetti et al., 2019, 2017), or together with complexes interconnecting them, such as Retromer or Retriever and the Commander complex, into which Retriever can be integrated (Bartuzi et al., 2016; Boesch et al., 2023a; Chandra et al., 2024; Healy et al., 2023; Mallam and Marcotte, 2017; McNally et al., 2017; Phillips-Krawczak et al., 2015; Seaman, 2021). Further coats, such as clathrin adapter complexes, VINE, and ACAP1-clathrin can also act on endosomes (Dai et al., 2004; Hooy et al., 2022; Li et al., 2007; Pang et al., 2014; Ren et al., 2013; Sachse et al., 2002; Shortill et al., 2022). These coat components organize a variety of endosomal protein exit routes with partially overlapping cargo specificity (Bean et al., 2017; Kvainickas et al., 2017; McGough et al., 2014; Steinberg et al., 2013), which may reflect their capacity to form hybrid carriers integrating different classes of sorting nexins (Gopaldass et al., 2026) .

Retromer is part of a protein coat mediating protein exit from endosomes in association with sorting nexins, such as Vps5, Vps17 or Snx3/Grd19 in yeast, and SNX27, ESCPE-1 or SNX3 in human cells (Seaman, 2021). Retromer consists of a VPS26–VPS29–VPS35 heterotrimer, which can form higher order complexes through VPS35 and VPS26 dimerization (Collins et al., 2008; Deatherage et al., 2020; Hierro et al., 2007; Kendall et al., 2022, 2020; Kovtun et al., 2018; Leneva et al., 2021; Lucas et al., 2016; Purushothaman et al., 2017). While sorting nexins themselves change the spontaneous curvature of membranes and can tubulate them (Kovtun et al., 2018; Leneva et al., 2021; Weering et al., 2012; Zhang et al., 2021), this activity can be enhanced by the association with Retromer, which may provide additional driving force for scaffolding the membrane (Gopaldass et al., 2023).

In human cells, numerous cargos recycle back to the cell surface in a Retromer-independent manner (Steinberg et al., 2013). Many of these depend on Retriever, an evolutionarily conserved complex of VPS26C/DSCR3, VPS29 and VPS35L/C16orf62, proteins that show distant homology to the Retromer subunits (McNally et al., 2017). Whereas the overall organization and shape of Retriever resemble those of Retromer, Retriever shows different cargo specificity and distinct features. Retriever forms part of a much larger complex, Commander, which integrates Retriever with the 12-subunit CCC complex and DENND10 and cooperates with the sorting nexin SNX17 to form coats (Boesch et al., 2023a; Butkovič et al., 2024; Healy et al., 2023; Martín-González et al., 2024; Singla et al., 2024; Steinberg et al., 2012). Like Retromer, also Commander can interact with WASH and thereby link to actin (Derivery et al., 2009; Gomez and Billadeau, 2009; Guo et al., 2024; Harbour et al., 2012; Jia et al., 2012; Phillips-Krawczak et al., 2015; Seaman et al., 2009).

Cargo carriers detach from endosomes by membrane fission (Gopaldass et al., 2024; Naslavsky and Caplan, 2023). Several factors have been implicated in this detachment, among them actin and the actin-controlling WASH complex (Derivery et al., 2009; Frisby et al., 2024; Gomez and Billadeau, 2009; Harbour et al., 2012; Jia et al., 2012; Seaman et al., 2009), dynamin-like GTPases and ATPases such as Vps1 and EHD1 (Arlt et al., 2015; Chi et al., 2014; Daumke et al., 2007; Deo et al., 2018; Gokool et al., 2007; Grant et al., 2001; Solinger et al., 2020), the ESCRT-III subunit IST1 (Clippinger et al., 2024) and the PROPPIN (β-propeller that binds phosphoinositides) WIPI1 and its yeast homolog Atg18 (DeLeo et al., 2021; Gopaldass et al., 2024, 2017). Coats and actin can exert forces on the underlying lipid bilayer, which not only allow the tubule to grow but may also produce friction between lipids and protein coats that contributes to fission (Johannes et al., 2014; Simunovic et al., 2018, 2017). In addition, dynamin-like proteins actively deform the bilayer to compress and destabilize it (Antonny et al., 2016). The lipid phosphatidylinositol-(3,5)-bisphosphate (PI(3,5)P_2_) and/or PI5P is also required for fission (DeLeo et al., 2021; Giridharan et al., 2022; Gopaldass et al., 2017; Rivero-Ríos and Weisman, 2022; Zieger and Mayer, 2012). PROPPINs are bona-fide effectors of PI(3,5)P_2_ (Courtellemont et al., 2022; DeLeo et al., 2021). They associate with membranes through two phosphoinositide binding sites (Baskaran et al., 2012; Krick et al., 2012; Liang et al., 2019; Proikas-Cezanne et al., 2004; Vicinanza et al., 2015; Watanabe et al., 2012). Upon membrane binding, a hydrophobic loop, which is located between the two lipid binding sites, folds into an amphipathic α-helix that inserts into the bilayer. Thereby, PROPPINs can tubulate lipid bilayers and drive their fission (Courtellemont et al., 2022; DeLeo et al., 2021; Gopaldass et al., 2017; Gopaldass and Mayer, 2024).

PROPPINs are a protein family that binds phosphatidylinositol 3-phosphate (PI3P), phosphatidylinositol 5-phosphate (PI5P) and phosphatidylinositol 3,5-bisphosphate (PI(3,5)P_2_) (Baskaran et al., 2012; Busse et al., 2015; Dove et al., 2004; Gopaldass and Mayer, 2024; Jeffries et al., 2004; Vicinanza et al., 2015). They are conserved from yeast (Atg18, Atg21, and Hsv2) to humans (WIPI1, WIPI2, WIPI3/WDR45B and WIPI4/WDR45) and exist in most eukaryotes in several isoforms, which can be grouped into two evolutionary clades (Dove et al., 2004; Proikas-Cezanne et al., 2015). In mammals, Atg18, Atg21, WIPI1 and WIPI2 belong to one clade, Hsv2, WIPI3 and WIPI4 to the second (Cong et al., 2021; Polson et al., 2010; Strong et al., 2021). WIPI1 and WIPI2 concentrate on phagophores when autophagy is induced and they are necessary for efficient formation of autophagosomes (Dooley et al., 2014; Polson et al., 2010; Proikas-Cezanne et al., 2004). WIPI2, which has a stronger impact on autophagy, interacts with different key factors for autophagosome formation.

WIPI1 and Atg18 also support membrane fission and protein sorting on endosomes and lysosomes, but for these functions they rely on molecular features that are distinct from those important to autophagy (Courtellemont et al., 2022; DeLeo et al., 2021; Gopaldass et al., 2017; Gopaldass and Mayer, 2024). WIPI1 localizes to endo-lysosomal compartments, where it affects the size of endosomes and the exit of proteins from endosomes to lysosomes (EGF-receptor), to the Golgi (Shiga-toxin), or to the cell surface (Transferrin receptor, GLUT-1) (DeLeo et al., 2021; Jeffries et al., 2004). Based on the differential effects of substitutions in the two phosphoinositide binding sites of WIPI1 it was suggested that binding of PI3P allows WIPI1 to support the formation of endosomal transport carriers while PI(3,5)P_2_ and lipid binding site two are needed for membrane fission at endosomes (DeLeo et al., 2021). WIPI1 as well as its yeast homolog Atg18 associate with Retromer to form the CROP complex, which amplifies the fission activity of the PROPPIN and makes its binding to membrane PI(3,5)P_2_ dependent (Courtellemont et al., 2022; Marquardt et al., 2023). This interaction depends on a conserved FSSS motif, which can be phosphorylated (Courtellemont et al., 2022; DeLeo et al., 2021; Feng et al., 2015; Gubas et al., 2024; Strong et al., 2021). This phosphorylatable FSSS motif exists also in WIPI2, where it forms also part of the binding site for ATG16L1, a core component of the autophagic machinery (Gubas et al., 2024; Strong et al., 2021) .

We have recently identified the Retromer- and WIPI1-containing CROP complex as an agent supporting the export of multiple cargos from endosomes, comprising both Retromer-dependent and Retromer-independent exit routes (Courtellemont et al., 2022; DeLeo et al., 2021). Since not all cargo exit from endosomes depends on WIPI1 and CROP we analysed alternative PROPPINs that might be involved in WIPI1-independent exit from endosomes. This led us to uncover a novel role for a WIPI2-Retriever complex in trafficking of β1-Integrin, which we present below.

## Results

WIPI1 promotes the exit of diverse cargos from endosomes, for example for transporting Transferrin to the plasma membrane, GLUT1 to the plasma membrane, EGF-receptor (EGFR) to lysosomes, and of Shiga toxin to the Golgi (DeLeo et al., 2021). β1-Integrin provides an exception, its delivery to the cell surface being unaffected by knockout of WIPI1 (Courtellemont et al., 2022). We investigated whether the endosomal exit of β1-Integrin might depend on another PROPPIN. Since WIPI1 and WIPI2 are similar (Bakula et al., 2013; Gopaldass and Mayer, 2024) belong to the same evolutionary clade, share two predicted lipid binding sites as well as the potential to form an amphipathic helix on their CD loop 6 (Cong et al., 2021; Gopaldass et al., 2017; Polson et al., 2010), we focused our attention on WIPI2B.

### WIPI1 and WIPI2 are specific for distinct trafficking pathways

We tested the impact of WIPI2 knockdown (WIPI2^KD^) on the afore-mentioned trafficking pathways. siRNA against WIPI2 reduced the protein in whole cell extracts by 94% when compared to cells transfected with control siRNA (Control) (Figure. S1). We analysed trafficking of EGFR from early endosomes to degradative, LAMP1-containing compartments. After 24h of serum starvation, cells were stimulated with EGF and analysed for EGFR localization by immunofluorescence. 15 min after EGF addition, EGFR-positive structures colocalized with the early endosomal marker EEA1 to more than 75% in both WIPI2^KD^ and control cells, indicating that EGF had induced efficient internalization of EGFR in both cases (Figure 1A-B). After 30 and 60 min of EGF stimulation, EGFR increasingly shifted from EEA1- towards LAMP1-positive lysosomal structures in control as well as in WIPI2-silenced cells. EGFR signals declined after 60 min in the immunofluorescence images as well as in Western blot analyses of the both cells (Figure. 1 C-E). Thus, WIPI2 knockdown does not impair EGFR trafficking from EEA1 endosomes to LAMP1-positive compartments and leaves lysosomal degradation of EGFR functional.

**Figure 1.**
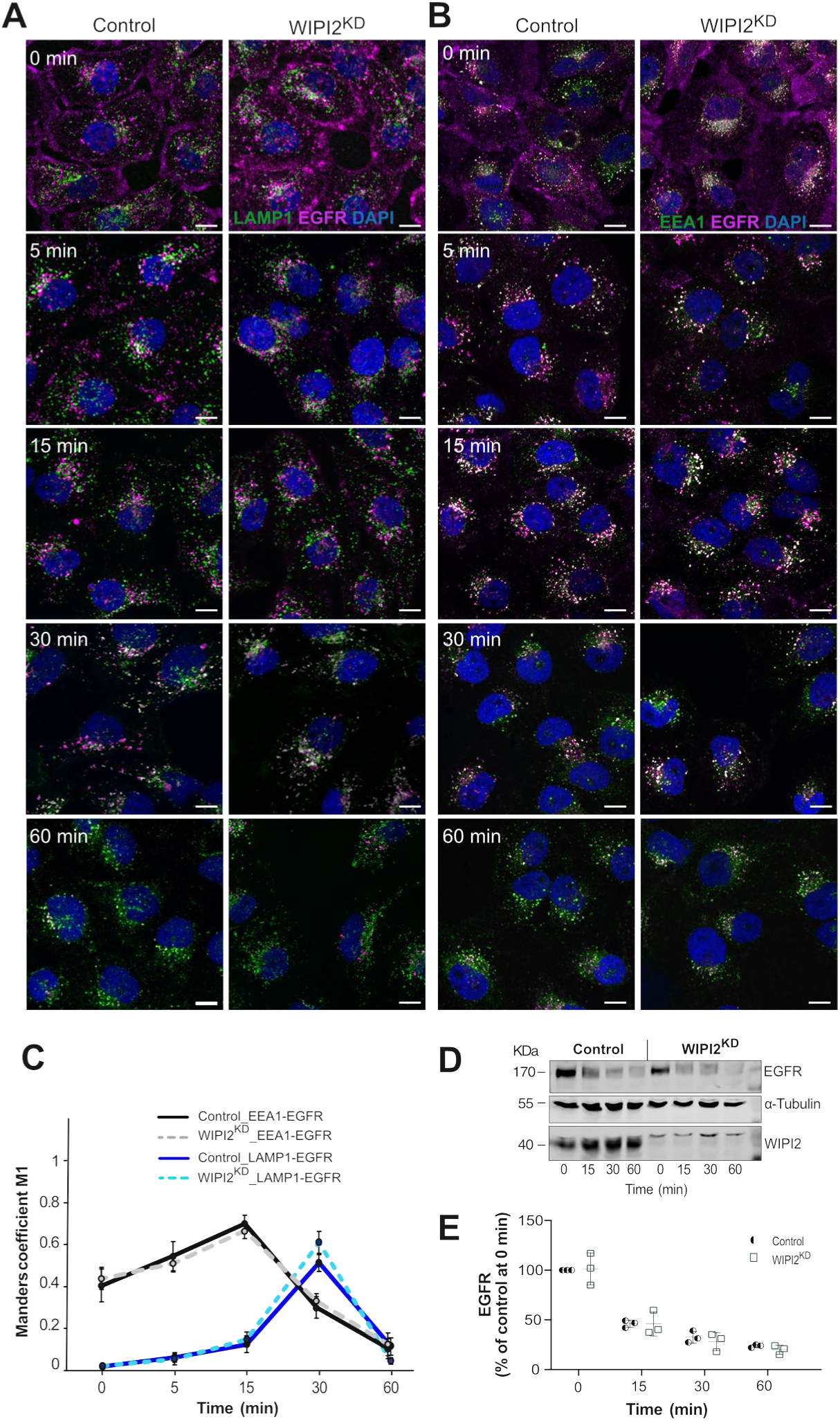
Effect of WIPI2 knockdown on EGFR degradation. **A. B.** Control and WIPI2^KD^ cells were serum-starved for 24 h and then supplemented with EGF (100 ng/ml). After the indicated periods of time, cells were fixed, permeabilized, DAPI-labeled (blue) and decorated with antibodies to EGFR (magenta), EEA1 or LAMP1 (green). Scale bars: 10 μm. **C.** Colocalization of EGFR with EEA1 or LAMP1 (white) was quantified over time using the images from A and B and Manders’ correlation coefficients were calculated. M1 indicates the fraction of magenta pixels overlapping with green pixels. Values are the mean ± s.d.; (n = 3). 150 cells, stemming from 3 independent biological experiments, were quantified per sample. **D.** EGFR degradation. Control and WIPI2^KD^ cells were stimulated with EGF for the indicated periods of time, lysed and subjected to SDS-PAGE and Western blot analysis for EGFR. α-Tubulin served as a loading control. **E.** Quantification of EGFR from (D), using the value of control cells at time 0 (cells starved for 24 h) as 100% reference. Data are means ±s. d., from three independent experiments. Data points form these three experiments are indicated for each time point.

To test the role of WIPI2 in Retromer-dependent trafficking we assayed the localisation of the glucose importer GLUT-1 because this protein accumulates on endosomes if its recycling to the plasma membrane through Retromer is impaired (Steinberg et al., 2013). To capture the surface fluorescence, we acquired z-stacks of non-permeabilized, immuno-stained cells, generated maximum intensity projections and integrated the fluorescence signal over the entire cell, normalized for its area as described in Materials and Methods section. Non-permeabilized WIPI2^KD^ cells showed similar cell surface staining of GLUT-1 as control cells (Figure 2 A-B), suggesting that also Retromer-mediated recycling of GLUT1 occurs independently of WIPI2.

**Figure 2:**
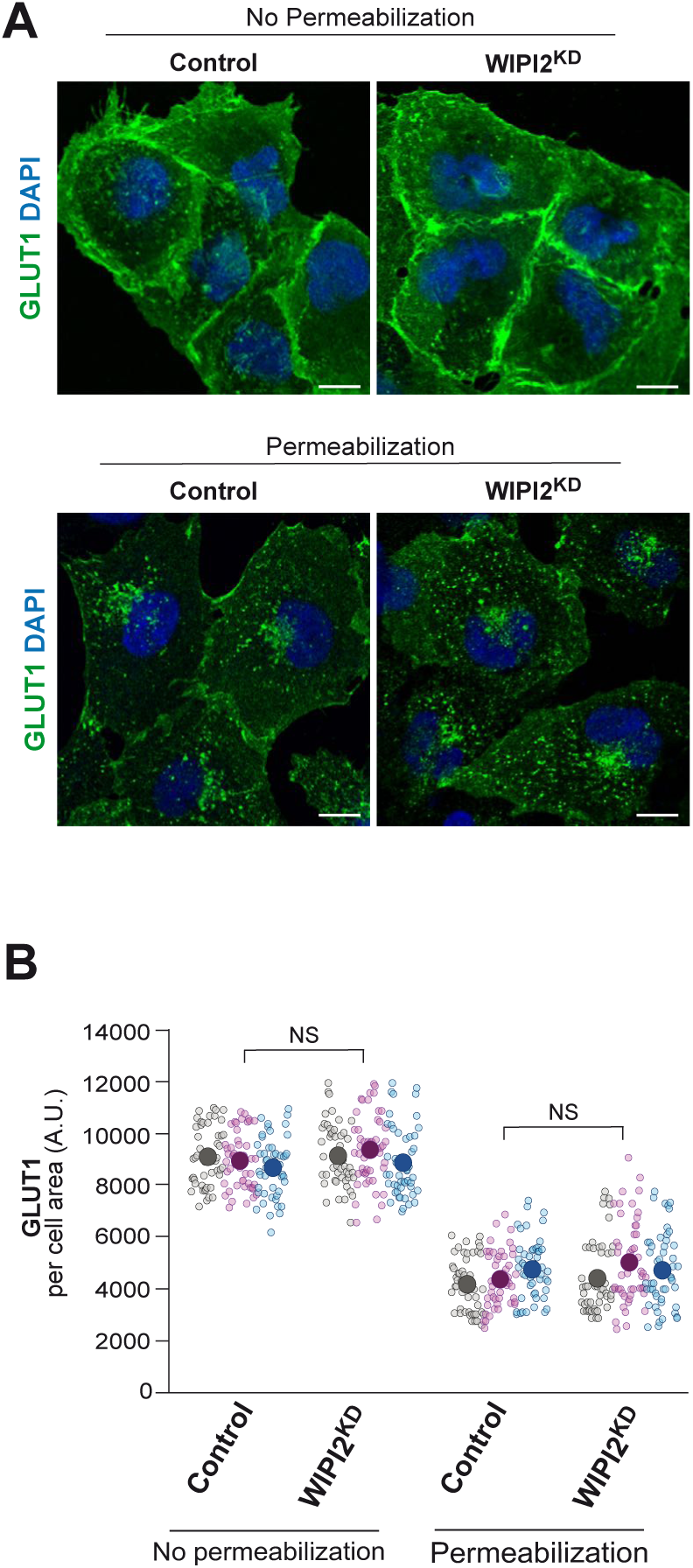
GLUT1 expression and localization upon WIPI2 knockdown. **A.** GLUT1 cell surface exposure. Control and WIPI2KD cells were fixed and stained with antibody against GLUT1 and with DAPI. Where indicated, cells had been permeabilized with 0.05% before staining to reduce plasma membrane staining and provide better access to GLUT1 inside the cell. Scale bars: 10 μm. **B.** Quantification of GLUT1-immunofluorescence in cells from A. Regions of interest (ROIs) corresponding to each cell and in some regions outside the cells (background) were manually defined using image J software. Total cell fluorescence was integrated, and the background fluorescence was subtracted. The resulting total cell fluorescence was divided by the area of the cell. This value is shown in the graph. 150 cells per condition, stemming from three independent experiments, were quantified. Values of individual cells and the means are presented by smaller and larger circles, respectively, coloured according to the independent experiment that generated them. P values were calculated applying an unpaired Student’s t-test with unequal variances. The analysis was performed with 99% confidence. NS = not significant (P > 0.05).

Next, we tested the requirement of WIPI2 for trafficking of β1-Integrin. Surface accumulation of this Integrin subunit is unperturbed by loss of SNX27 or Retromer but depends on the SNX17-Retriever pathway (McNally et al., 2017; Steinberg et al., 2013). Knockdown of WIPI2 induced a loss of β1-Integrin immunofluorescence from the surface of HK2 cells (Figure 3 A, B) and phenocopied the previously published knockdown of SNX17 (McNally et al., 2017; Steinberg et al., 2013). When the cells were detergent-permeabilized to partially remove the plasma membrane and improve accessibility of internal structures for immunostaining, cells depleted of WIPI2 showed an accumulation of β1-Integrin on intracellular structures that colocalized with the early endosomal marker EEA1 and, to a lesser degree, with the late endosomal/lysosomal marker LAMP1 (Figure 3 C-E,G). WIPI2 knockdown had no impact on the total EEA1 and LAMP1 fluorescence level per cell, suggesting that the abundance of these compartments remained the same (Figure 3 F,H). In accord with previous observations (DeLeo et al., 2021), β1-Integrin surface labelling in non-permeabilized cells was not affected by knockdown of WIPI1 (Figure S2). Thus, β1-Integrin recycling to the plasma membrane selectively requires WIPI2.

**Figure 3.**
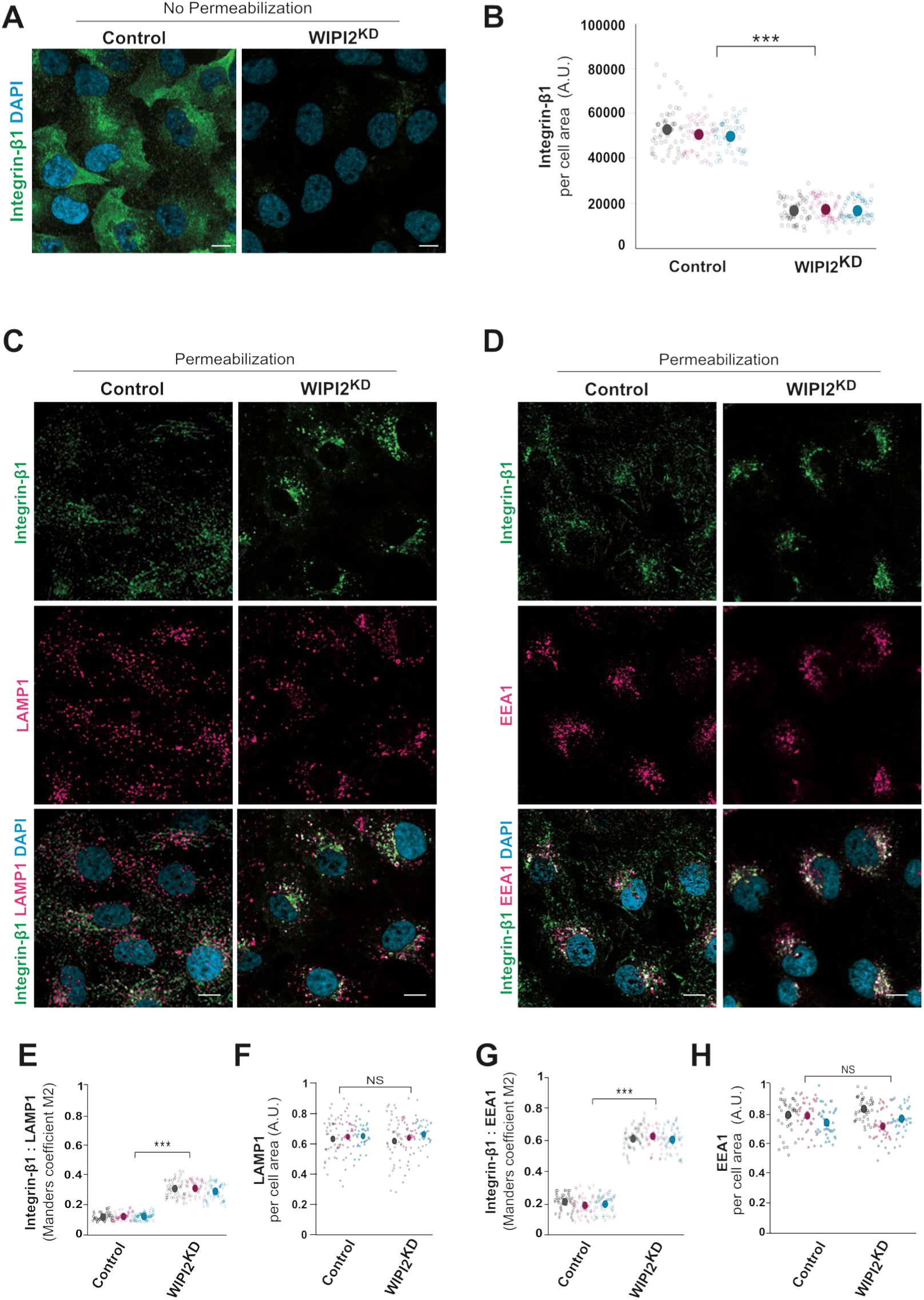
Impact of WIPI2 knockdown on β1-Integrin localization. **A.** β1-Integrin cell surface expression. Control and WIPI2^KD^ cells were fixed and stained with antibody to β1-Integrin (green) and with DAPI (blue), without detergent permeabilization. Scale bars: 10 μm. **B.** Quantification of β1-Integrin immunofluorescence in cells from A. Regions of interest (ROIs) corresponding to each cell and to areas outside the cells (background) were manually defined using ImageJ software. Total cell fluorescence was integrated, and the background fluorescence was subtracted. The resulting total cell fluorescence was divided by the area of the cell. This value is shown in the graph. 150 cells per condition, stemming from three independent biological experiments, were quantified. Individual cells and means are presented by smaller and larger circles, respectively, coloured according to the independent experiment that they stem from. P values were calculated through an unpaired Student’s t-test with unequal variances.The analysis was performed with 99% confidence. ***P < 0.001. **C.D.** Immunofluorescence staining of intracellular β1-Integrin (green) and EEA1 (magenta) or LAMP1 (magenta) in control and WIPI2^KD^ cells. Overlaps are marked in white. Cells had been fixed, permeabilized with 0.05% saponin and stained with antibodies to the indicated proteins. Scale bars: 10 μm. **E.G.** Colocalization of β1-Integrin with LAMP1 or EEA1 was measured in cells from C, using Manders’ colocalization coefficient M2, calculated in Image J software. It indicates the fraction of green pixels overlapping with the magenta pixels. Colocalization was quantified from three independent experiments with a total of 150 cells per condition. Values of individual cells and means are presented by smaller and larger circles, respectively, coloured according to the independent experiment that they stem from. An unpaired Student’s t-test with unequal variances was used to calculate P-values. The analysis was performed with 99% confidence. ***P < 0.001. **F.H.** Quantification of LAMP1- or EEA1-immunofluorescence in cells from C or D, respectively, was performed as in B. 120 cells per condition, stemming from three independent biological experiments, were quantified. Individual cells and means are presented by smaller and larger circles, respectively, coloured according to the independent experiment that they stem from. Data are means ± s.d. P-values were calculated applying an unpaired Student’s t-test with unequal variances. The analysis was performed with 99% confidence. NS = not significant (P > 0.05).

### The conserved amphipathic α-helical and FSSS motifs of WIPI2 promote β1-Integrin recycling

WIPI2 shares two critical features with other PROPPINs (Gopaldass and Mayer, 2024). These include an FSSS motif and the CD loop on blade 6 of the β-propeller structure. The CD loop promotes membrane fission activity of Atg18 during the fragmentation of yeast vacuoles (Gopaldass et al., 2017) and in the detachment of endosomal carriers (DeLeo et al., 2021). Upon contact with the bilayer, the loop folds into an amphipathic α-helix that is conserved among PROPPINs and necessary for membrane fission activity. To test whether the amphipathic nature of this helix also contributes to the function of WIPI2 in β1-Integrin retrieval to the cell surface, we swapped two pairs of amino acids on opposite sides of this α-helix in WIPI2 (Figure 4 A). This manipulation preserves the amino acid composition of the helix but abolishes its amphipathic character. The resulting ^EGFP^WIPI2^SLoop^ (scrambled loop) variant and the wildtype version ^EGFP^WIPI2^WT^ were expressed in control and WIPI2^KD^ cells at comparable levels, as shown by Western blot (Figure S3). Whereas permeabilized WIPI2^KD^ cells displayed β1-Integrin concentrated on perinuclear endosomes, expression of ^EGFP^WIPI2^WT^ rescued this phenotype (Figure 4 B). ^EGFP^WIPI2^Sloop^ failed to induce such recovery in WIPI2^KD^ cells. It even exerted a dominant-negative effect when expressed in control cells. (Figure 4 C-D). Cells expressing ^EGFP^WIPI2^SLoop^ also showed conspicuous long tubules, which were more readily detectable in live cell imaging (Figure S4) than in fixed cells (Figure 4).

**Figure 4.**
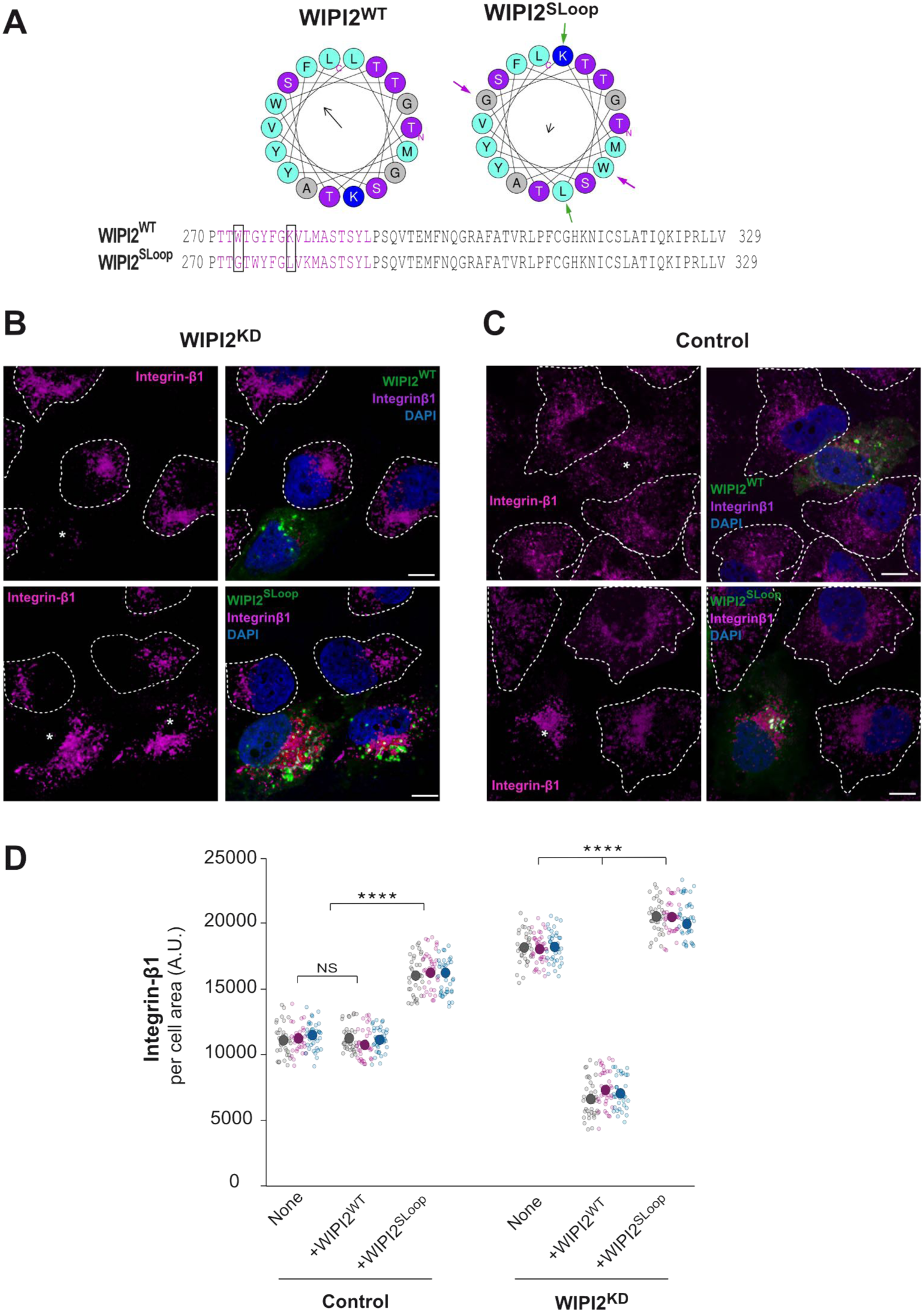
Role of the amphipathic α-helix of WIPI2 in β1-Integrin sorting. **A.** The amphipathic α-helix. Helical wheel projections showing the CD-loop on blade 6 of the wildtype sequence, WIPI2^WT^, and WIPI2^Sloop^. Coloured arrows indicate the two pairs of amino acids that have been swapped in WIPI2^Sloop^. The magnitude and direction of the hydrophobic moment of the helices was predicted using the online tool Heliquest (Gautier et al., 2008). It is indicated by the vector in the centre of the wheels. Sequences of the hydrophobic loop region of WIPI2^WT^ and WIPI2^Sloop^ are shown, and predicted α-helices are plotted in magenta. The two pairs of hydrophobic/hydrophilic amino acids that are swapped in WIPI2^Sloop^ are highlighted by rectangles in the sequences. **B.C.** β1-Integrin localisation. WIPI2^KD^ (B) and control (C) cells were transfected with a plasmid carrying siRNA-resistant ^EGFP^WIPI2^WT^ or ^EGFP^WIPI2^Sloop^. After 18h of viral transfection, cells were fixed, permeabilized with 0.05% saponin and stained with DAPI and antibodies to β1-Integrin. Dashed lines indicate the circumference of untransfected cells, while transfected cells are indicated by asterisks. Scale bars: 10 μm. **D.** Quantification of β1-Integrin immunofluorescence in cells from B and C. Regions of interest (ROIs) corresponding to cells expressing the indicated WIPI2 variants, and some regions outside the cells (background), were manually defined using Image J software. Total cell fluorescence was integrated and the background fluorescence was subtracted. The resulting total cell fluorescence was divided by the area of the cell. This value is shown in the graph. 105 cells per condition, stemming from three independent biological experiments, were quantified. Values of individual cells and means are presented by smaller and larger circles, respectively, coloured according to the independent experiment that generated them. P-values were calculated applying an unpaired Student’s t-test with unequal variances. The analysis was performed with 99% confidence: ****P < 0.00001. NS: not significant (P>0.05)

Alignments of Atg18 and WIPI1/2 orthologs have revealed a conserved FSS/TS motif in blade 2 (Courtellemont et al., 2022; Gopaldass et al., 2017; Gopaldass and Mayer, 2024; Strong et al., 2021). On the β-propeller structure of these PROPPINS, this motif is located at the membrane-distal side. It is a potential phosphorylation site and required for the interaction of Retromer with WIPI1 and Atg18 (Courtellemont et al., 2022; Feng et al., 2015). Since the overall structural arrangement of Retriever resembles that of Retromer (Bartuzi et al., 2016; Boesch et al., 2023a; Healy et al., 2023; McNally et al., 2017), we tested whether WIPI2 requires its FSSS motif to bind Retriever. To this end, we generated WIPI2^S67E^ and WIPI2^S67A^ to substitute the central serine in the FSSS motif and mimic effects of potential phosphorylation at this site.

The functionality of these WIPI2 variants was tested by rescuing WIPI2^KD^ cells through expression of siRNA-resistant variants of ^EGFP^WIPI2. Both ^EGFP^WIPI2^S67E^ and ^EGFP^WIPI2^S67A^ were expressed to similar levels as an ^EGFP^WIPI2 wildtype allele (Figure S3). Immunofluorescence staining of non-permeabilized WIPI2^KD^ cells showed that the expression of ^EGFP^WIPI2^WT^ re-established the exposure of β1-Integrin at the cell surface, while ^EGFP^WIPI2^S67A^ and ^EGFP^WIPI2^S67E^ were ineffective (Figure 5 A-C). Expression of the two variants also impaired surface exposure of β1-Integrin in control cells (Figure 5 B-D). This dominant negative effect is significant because it implies that the two variants compete with the endogenous protein. To be able to do so they should be folded, suggesting that the S67 substitutions do not grossly perturb WIPI2 structure. A conspicuous consequence of both FSSS substitutions was the accumulation of ^EGFP^WIPI2^S67A^ and ^EGFP^WIPI2^S67E^ in long tubules. These tubules were very extensive in live cell imaging and easily decayed upon fixation. (Figures. 5 and S4). Co-expression of ^mCherry^RAB4 or ^mCherry^Rab5 revealed that these tubules often connected larger dot-like structures, which were positive for ^mCherry^Rab5 and/or ^mCherry^RAB4 (Figure S4 A). The tubules themselves preferentially accumulated ^mCherry^RAB4 (70% colocalization) and only little ^mCherry^Rab5 (21%) (Figure S4 B). These results are consistent with the view that WIPI2-associated tubules originate mainly from early and recycling endosomes (Figure S4) and that WIPI2 contributes fission activity to Retriever-mediated trafficking through its amphipathic helix and its FSSS motif.

**Figure 5.**
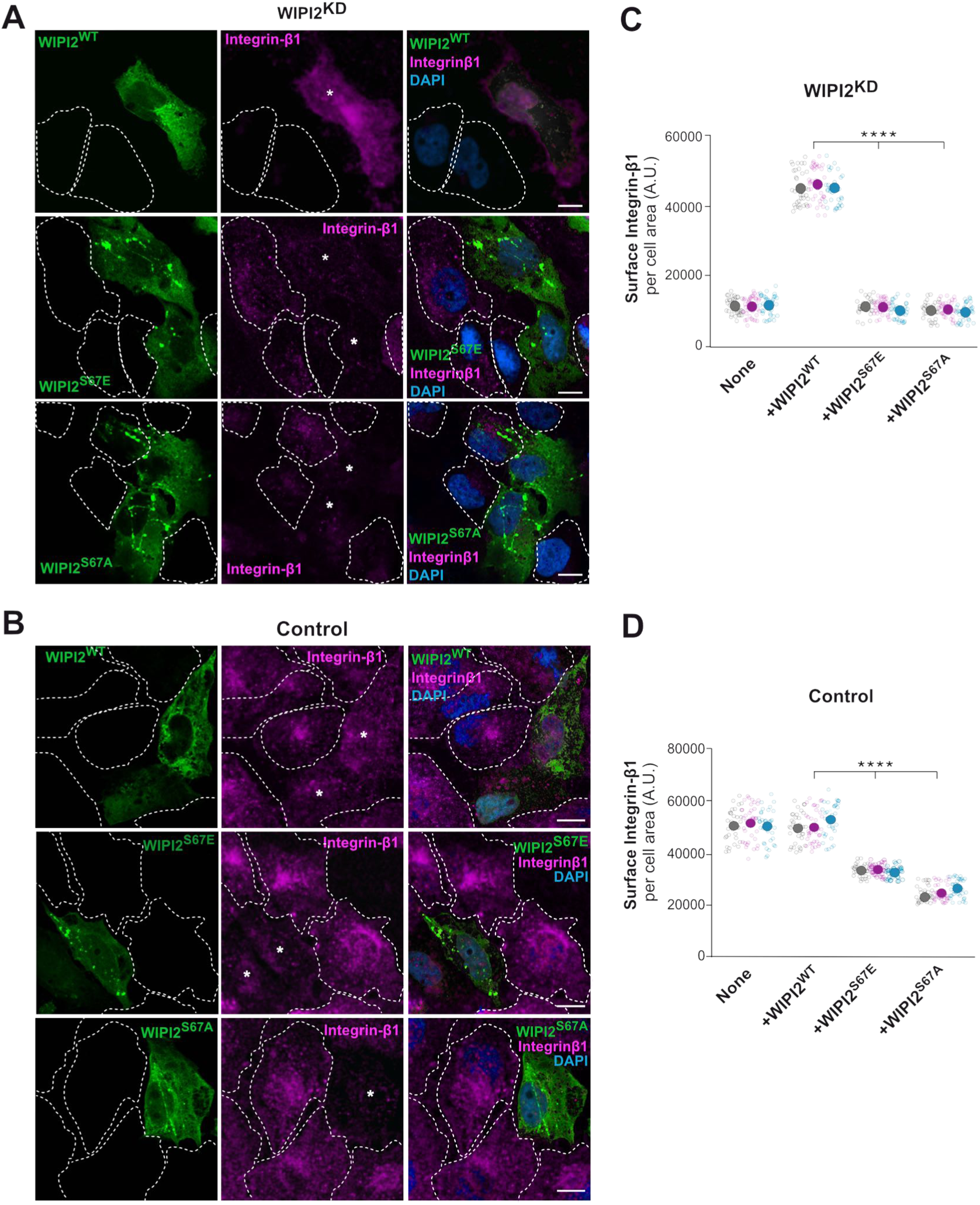
Effects of the WIPI2 FSSS motif on β1-Integrin recycling. **A.B.** Influence of WIPI2 variants on β1-Integrin. WIPI2-knockdown (WIPI2^KD^, A) and control (B) HK2 cells were transfected with a plasmid expressing siRNA-resistant wildtype or the indicated FSSS variants of ^EGFP^WIPI2. After 18 h of viral transfection, cells were fixed (without detergent permeabilization) and stained with DAPI and antibodies to β1-Integrin (red). Dashed lines indicate the circumference of untransfected cells, while transfected cells are indicated by asterisks. Scale bars: 10 μm. **C. D .** Quantification of β1-Integrin immunofluorescence in cells from A and B was performed as in Figure 4 D. 105 cells per condition, stemming from three independent biological experiments, were quantified. Individual cells and means are presented by smaller and larger circles, respectively, coloured according to the independent experiment that they stem from. P-values were calculated applying an unpaired Student’s t-test with unequal variances. The analysis was performed with 99% confidence: ****P < 0.00001.

### WIPI2 interacts with Retriever, Commander and SNX17

Next, we tested whether WIPI2 and WIPI1 interact with Retriever or Retromer and could become part of their respective coats. In that case, WIPI2 should interact not only with Retriever itself, but also with the sorting nexin SNX17, which binds Retriever, and perhaps the CCC subunits, which integrate Retriever into the Commander complex (Butkovič et al., 2024; Martín-González et al., 2024; Singla et al., 2024). We assayed this by co-immunoadsorption. An HA-tagged allele (WIPI2^HA^) was expressed in HK2 cells using lentiviruses. Extracted proteins were adsorbed to anti-HA beads and analysed by Western blotting (Figure 6). Whereas the Commander subunit CCDC93 and the Retriever subunit VPS26C co-adsorbed to the beads in extracts from cells expressing WIPI2^HA^ (Figure 6 A, C), these signals were absent from cells expressing only non-tagged WIPI2. SNX17 also co-adsorbed with WIPI2^HA^. This interaction was abolished by knockdown of the Retriever subunit VPS26C (Figure 6 B), suggesting that Retriever links WIPI2 to SNX17. WIPI2 differentiated strongly between Retromer and Retriever. Although VPS26C is quite similar to the Retromer subunit VPS26, only VPS26C co-adsorbed with WIPI2^HA^ whereas VPS26 was not detectable (Figure 6 C, D).

**Figure. 6:**
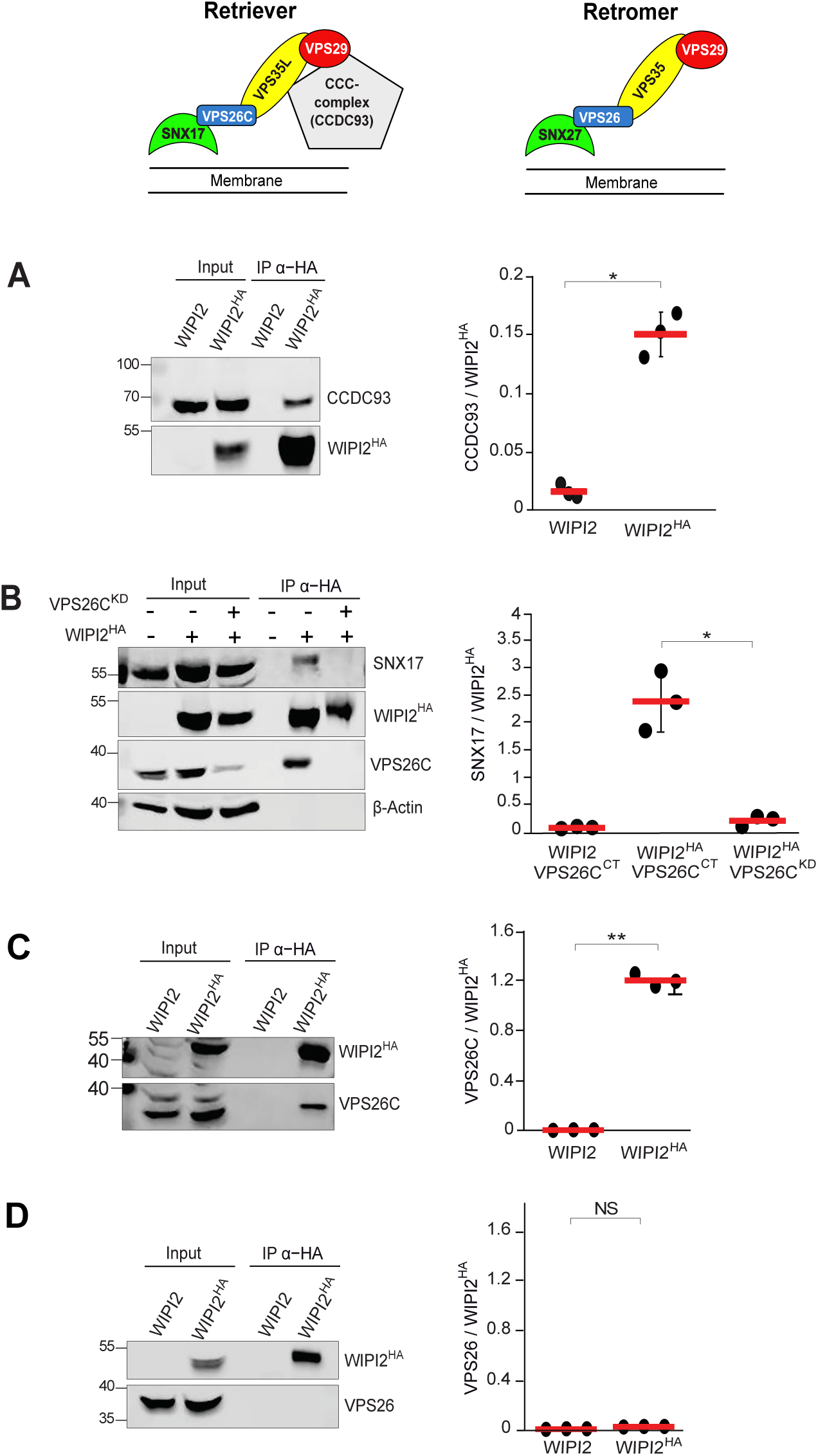
Integration of WIPI2 with coat subunits. **A.** Interaction of WIPI2^HA^ with the CCC complex subunit CCDC93. HK2 cells and HK2 cells stably expressing WIPI2^HA^ were detergent solubilised, the total cell extracts were incubated with anti-HA beads and washed. Adsorbed protein was analysed by SDS-PAGE and Western blotting using the indicated antibodies. The intensity of the interacting CCDC93 was quantified with a LICOR Odyssey fluorescence imager. The background from the corresponding position in the sample from cells without HA-tag was subtracted. The resulting intensity is shown relative to the intensity of WIPI2^HA^ signal on the beads. N = 3 biological replicates from independent experiments. Red bars show the means. Error bars represent the SEM. P values were calculated applying an unpaired Student’s t-test with unequal variances. *P < 0.01. **B.** Retriever-dependent interaction of WIPI2^HA^ and SNX17. HK2 cells stably expressing WIPI2^HA^ were treated with siRNA to VPS26C (VPS26C^KD^), or with non-specific siRNA (VPS26C^CT^), and lysed. The total cell extracts were incubated with anti-HA beads and adsorbed proteins were analysed using the indicated antibodies as in A. Values shown stem from 3 independent experiments. **C. D.** Selectivity for Retriever versus Retromer. The co-immunoadsorption experiments were performed as in A and analysed for co-adsorbed (C) VPS26C (Retriever) or (D) VPS26 (Retromer). N=3 independent experiments. *P < 0.01.**P < 0.001. NS=not significant.

The situation was inversed when the HA tag was attached to WIPI1. In this case, VPS26 was recovered on the beads but VPS26C was not (Figure 7A, B). SNX27, a sorting nexin that interacts with Retromer (Simonetti et al., 2022; Temkin et al., 2011), was also co-adsorbed with WIPI1^HA^ (Figure 7C). The immunoprecipitation results were supported by parallel immunofluorescence studies. WIPI2 colocalized preferentially with Retriever and SNX17 (Figure S5) while WIPI1 colocalized preferentially with SNX27 and Retromer (Figure S6). Moreover, WIPI2 colocalized with integrin-β1 on early endosomal compartments positive also for SNX17, but not on late endosomal LAMP1 containing compartments (Figure S7). These results suggest that Retriever bridges WIPI2 and SNX17 and that Retromer bridges WIPI1 with SNX27, integrating these PROPPINs into their cognate Retriever- or Retromer- coats.

**Figure. 7:**
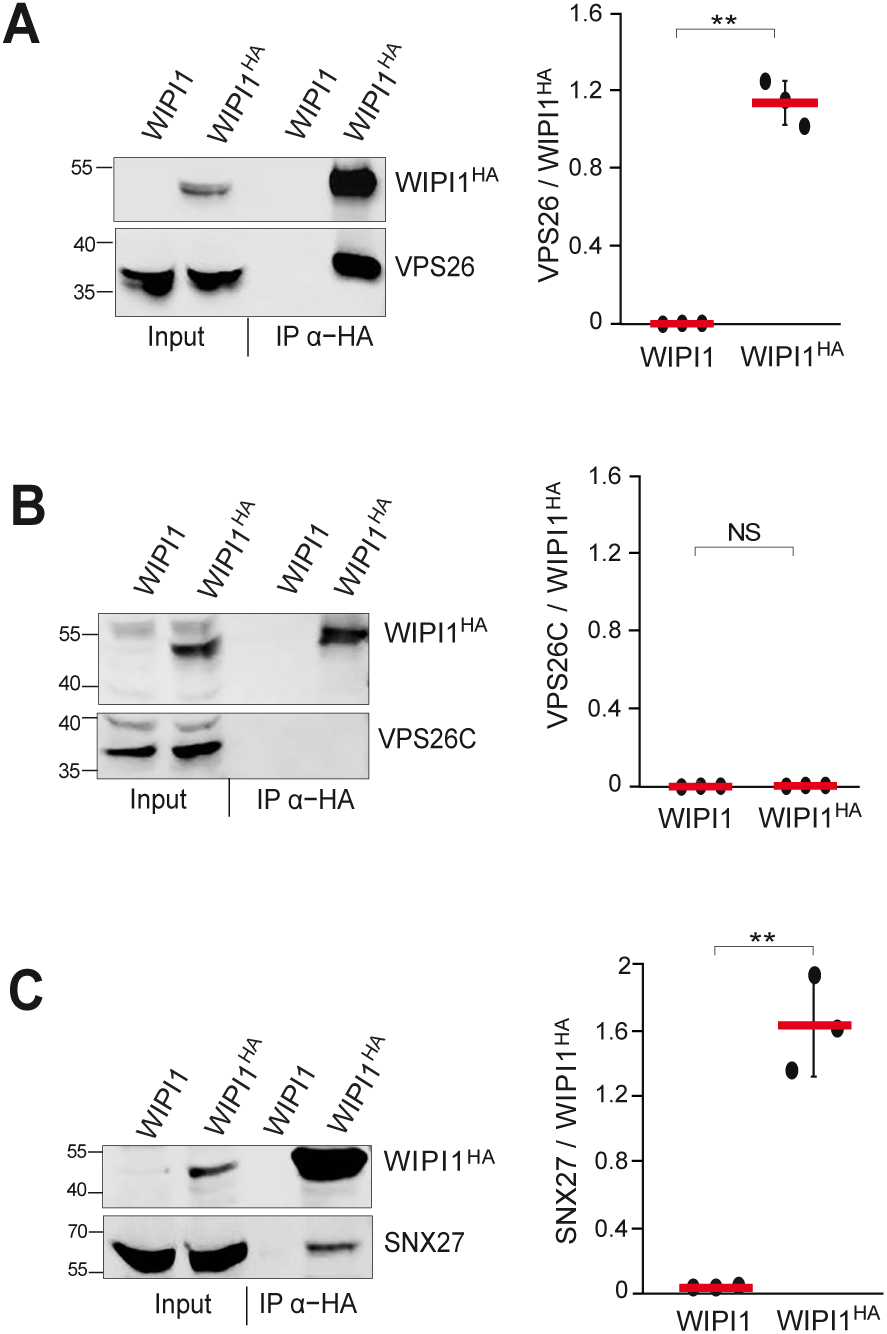
Interaction of WIPI1 with VPS26 and SNX27, but not VPS26C. **A.** Interaction of WIPI1^HA^ with VPS26. HK2 cells and HK2 cells stably expressing WIPI1^HA^ were detergent-solubilised, the total cell extracts were incubated with anti-HA beads and washed. Adsorbed protein was analysed by SDS-PAGE and Western blotting using the indicated antibodies. The intensity of the interacting VPS26 was quantified with a LICOR Odyssey fluorescence imager. The background from the corresponding position in the sample from cells without HA-tag was subtracted. The resulting intensity is shown relative to the intensity of the WIPI1^HA^ signal on the beads. N = 3 biological replicates from independent experiments. Red bars show the means and error bars represent the SEM. P values were calculated applying an unpaired Student’s t-test with unequal variances. **B.** Lack of interaction of WIPI1^HA^ with VPS26C. HK2 cells stably expressing WIPI1^HA^ were used for co-immunoadsorption experiments as in A and decorated with the indicated antibodies. N = 3. **C.** Interaction of WIPI1^HA^ with SNX27. HK2 cells stably expressing WIPI1^HA^ were used for co-immunoadsorption experiments as in A and decorated with the indicated antibodies. N = 3. **P < 0.001. NS: not significant.

Since the FSSS motif of WIPI2 is necessary for its transport functions, we tested whether this motif affects the interactions of WIPI2 with Retriever. To this end, wildtype and the two FSSS variants of WIPI2^HA^ were stably expressed and cell extracts were probed by the same co-adsorption approach as above (Figure 8). Whereas the Retriever subunit VPS26C was efficiently co-adsorbed with WIPI2^WT-HA^, WIPI2^S67A-HA^ and WIPI2^S67E-HA^ labilized this interaction. Cells carrying only the non-tagged, endogenous WIPI2 yielded no VPS26C signal. Thus, the FSSS motif is required for the interaction between WIPI2 and Retriever. These results were also confirmed by immunofluorescence analysis (Figure S8), where VPS26C colocalized less with WIPI2^S67A^ and WIPI2^S67E^ (16%) than with the wildtype form (56%).

**Figure. 8:**
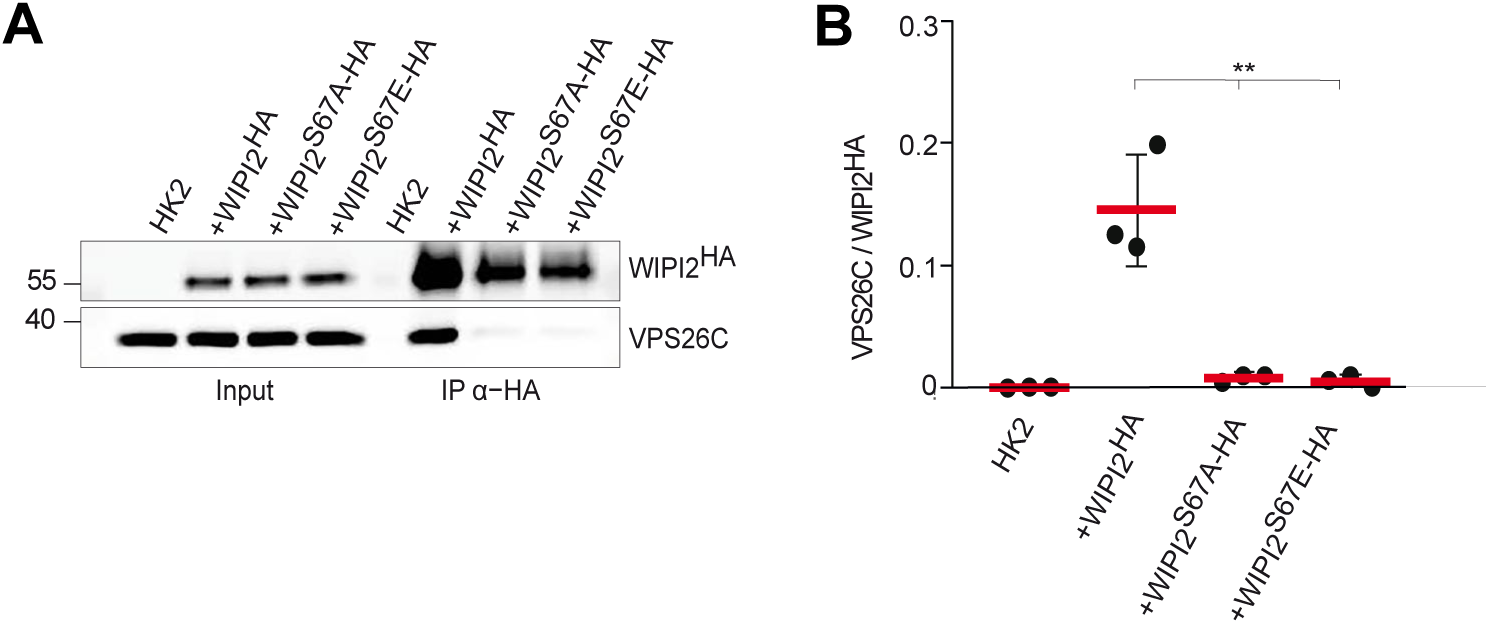
Impact of the FSSS motif on the WIPI2-Retriever interaction. **A.** HK2 cells and HK2 cells stably expressing the indicated WIPI2^HA^ variants were detergent solubilised. Anti-HA beads were incubated with the total cell extracts, washed, and adsorbed proteins were analysed by SDS-PAGE and Western blotting using the indicated antibodies. **B.** Band intensities from the blots in A were quantified with a LICOR Odyssey infrared fluorescence imager and plotted as the ratio of VPS26C over WIPI2^HA^. N=3 independent experiments were quantified. Red bars show the means and error bars represent the SEM. P values were calculated applying an unpaired Student’s t-test with unequal variances. **P < 0.001.

### The interaction of WIPI2 and Retriever is necessary for their localization on recycling endosomes

To test for the in vivo consequences of the WIPI2-Retriever interaction, we analysed the effect of the FSSS motif on the colocalization of the respective WIPI2^EGFP^ variants with different endosomal RAB-GTPases, which were N-terminally tagged with mCherry. In line with previous studies (Polson et al., 2010; Puri et al., 2018) ^EGFP^WIPI2^WT^ mostly co-localized with ^mCherry^RAB11 (58%) and ^mCherry^RAB5 (44%) and much less with and ^mCherry^RAB7 (6%) (Figures 9 and S9-S11). By contrast, ^EGFP^WIPI2^S67A^ and ^EGFP^WIPI2^S67E^ co-localized mainly with ^mCherry^RAB5 (60-63%) and ^mCherry^RAB7 (36-38%), and much less with ^mCherry^RAB11 (12-14%). The Retriever subunit VPS35L was detected by immuno-staining. In cells expressing WIPI2^WT^, it mainly co-localized with ^mCherry^RAB11 (42%) and ^mCherry^RAB5 (43%) and less with ^mCherry^RAB7 (23%) (Figures 9 and S9-S11). In ^EGFP^WIPI2^S67E^ and ^EGFP^WIPI2^S67A^ expressing cells, VPS35L distribution shifted, similarly as for ^EGFP^WIPI2, indicating reduced presence on ^mCherry^RAB11 compartments (12-13%), whereas co-localisation with ^mCherry^RAB5 (42-44%) was maintained and that with ^mCherry^RAB7 (26-27%) slightly increased. Collectively, these results suggest that the FSSS motif in blade 2 of WIPI2 allows this PROPPIN to associate with Retriever, and that this association is necessary for localising WIPI2 and Retriever on Rab11 containing recycling endosmes.

**Figure 9.**
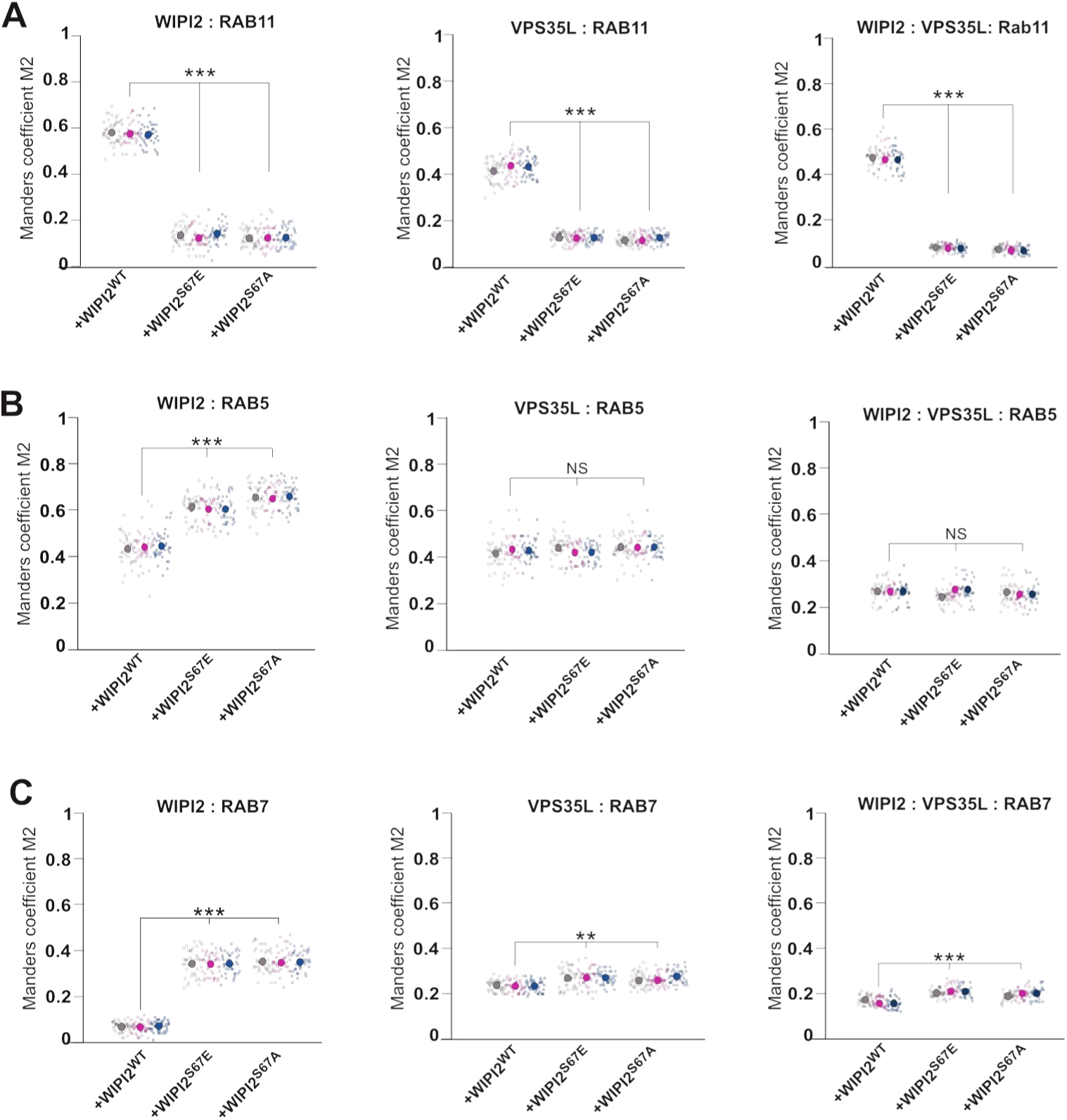
Role of the WIPI2 FSSS motif for recruiting WIPI2 and Retriever to Rab11 endosomes. WIPI2-knockdown HK2 cells were transfected with plasmids expressing the indicated siRNA-resistant ^EGFP^WIPI2 variants and mCherry-RAB5, mCherry-RAB7, or DsRed-Rab11. Cells were fixed, permeabilized, immuno-stained for VPS35L and analyzed by confocal microscopy. Examples of the quantified images are presented in Supplementary Figures S9 to S11. Colocalization with ^EGFP^WIPI2 and VPS35L was assessed for: **A.** RAB11 **B.** RAB5 **C.** RAB7. Colocalization was quantified using Manders’ colocalization coefficient M2, calculated in ImageJ. M2 refers to the fraction of VPS35L colocalising with the different RAB-proteins. For the triple colocalizations WIPI2/VPS35L/RAB, M2 indicates the fraction of green pixels (WIPI2) overlapping with the pixels positive for the VPS35L/RAB colocalization. Colocalization was quantified from three independent biological experiments in a total of 120 cells. Individual cells. Individual cells and means are presented by smaller and larger circles, respectively, coloured according to the independent experiment that they stem from. P-values were calculated by an unpaired Student’s t-test with unequal variances. The analysis was performed with 99% confidence. **P < 0.001; ***P < 0.0001. NS = not significant (P>0.05).

## Discussion

Our experiments reveal pathway specificity for WIPI1 and WIPI2. The two proteins support distinct cargoes and transport routes and interact with different partners.

This isoform selectivity contrasts with the roles of these proteins in their second functional domain, autophagy, which strongly depends on WIPI2 but is also significantly reduced by ablation of WIPI1 (DeLeo et al., 2021; Dooley et al., 2014; Guan et al., 2001; Obara et al., 2008; Polson et al., 2010; Proikas-Cezanne et al., 2015, 2004). Furthermore, the autophagic function of the PROPPINs also exploits other molecular features than endosomal protein exit. For autophagy, WIPI1 and its yeast orthologue Atg18 do not require the amphipathic helix in CD loop 6, which is essential for endosomal transport. It was hence proposed that WIPI1 and Atg18 perform distinct functions in autophagy and in endosomal protein exit (Courtellemont et al., 2022; DeLeo et al., 2021; Gopaldass et al., 2017).

In endosomal transport, WIPI1 and WIPI2 display selectivity in their partners (Fig. 10). WIPI1 interacts with Retromer and WIPI2 with Retriever, mirroring the impact of the two PROPPINs on the respective transport routes and cargos. WIPI1 and WIPI2 utilise the same molecular features to support these transport routes. Their FSSS motif is necessary for both isoforms to integrate with Retromer and Retriever, respectively. This, together with the similarity in the overall organisation of Retromer and Retriever, suggest that they might form complexes in a similar manner. In analogy to the WIPI1-Retromer complex, which was termed CROP (Courtellemont et al., 2022), we hence refer to the WIPI2-Retriever complex as CROP2.

WIPI2-dependent transport of β1-Integrin depends on the amphipathic character of the α-helix in CD loop 6. In Atg18 and WIPI1, this helix is necessary to convey membrane fission activity, which is used both for division of the lysosome-like vacuoles of yeast and for detachment of endosomal carriers (DeLeo et al., 2021; Gopaldass et al., 2017; Zieger and Mayer, 2012). This fission activity is potentiated by integrating Atg18 and WIPI1 with Retromer into the CROP complex (Courtellemont et al., 2022). Our observations suggest that WIPI2 contributes pathway-specific membrane fission activity by integrating with Retriever. In line with this, ablating the amphipathic character of its helix on CD loop 6 disrupts β1-Integrin recycling to the cell surface and leads to its accumulation in endosomal compartments. Disrupting the WIPI2-Retriver interaction by substitution of the FSSS motif has the same effect. It also leads to the accumulation of long tubular structures, which we presume to be exaggerated endosomal carriers that continue to grow but fail to detach. This phenotype is consistent with a block in the membrane fission step that is necessary to allow the endosomal carrier to depart.

Retriever couples to other proteins that are required for endosomal protein transport. These include the CCC complex, which associates with Retriever to yield the very large Commander complex (Boesch et al., 2023b; Healy et al., 2023; Laulumaa et al., 2024). How Commander is placed on the membrane is currently unknown, but the interaction between SNX17 and Retriever contributes to its recruitment (Butkovič et al., 2024; Healy et al., 2022; Singla et al., 2024; Steinberg et al., 2012) and presumably to the formation of a coat. Our pull-downs of WIPI2 detected not only the Retriever subunit VPS26C as an interactor, but also SNX17 and the CCC subunit CCDC93. This co-fractionation could be due to a direct interaction of WIPI2 with Retriever or reflect an interaction via another component of Commander. Our immunoprecipitation assays cannot distinguish and more detailed structural and interaction studies with pure compounds will be necessary to elucidate the nature of this interaction. Since Retriever is structurally similar to Retromer, WIPI2 is structurally similar to WIPI1 and Atg18, and a Retromer-Atg18 interaction could be reconstituted from pure proteins (Courtellemont et al., 2022) , we consider a direct WIPI2-Retriever interaction as the more likely scenario. But in both cases our observations suggest that WIPI2 integrates into the Retriever coat and might thus bring membrane fission activity to the place where it is required for carrier formation. Here, it could synergize with other factors that support membrane fission in endosomal transport, such as EHD1, which interacts with SNX17, and the actin-regulating WASH complex, which binds to Commander (Bartuzi et al., 2016; Dhawan et al., 2022, 2020; Phillips-Krawczak et al., 2015).

The requirement of two distinct WIPI proteins, using the same molecular features, in Retromer- and Retriever-dependent transport uncovers a novel conserved element between these two pathways. In combination with the similar overall organisation of Retromer and Retriever and their common interactions with EHD proteins (Arlt et al., 2015; Chi et al., 2014; Daumke et al., 2007; Deo et al., 2018; Gokool et al., 2007; Grant et al., 2001; Solinger et al., 2020) and WASH (Derivery et al., 2009; Gomez and Billadeau, 2009; Guo et al., 2024; Harbour et al., 2012; Jia et al., 2012; Phillips-Krawczak et al., 2015; Seaman et al., 2009), this supports the notion that these two membrane coats share similar mechanistic principles for their formation and/or detachment.

## Materials and Methods

### Antibodies and reagents

All chemical reagents were from Sigma-Aldrich unless specified otherwise. We used anti-LAMP1 (H4A3, US Biological, Life Sciences); anti- EEA1 (#2411, Cell Signaling); anti-EGFR (sc-373746, clone A-10 Santa Cruz Biotechnology; PA5-85089 ThermoFisher Scientific); anti-Integrin β1 ( ab150361, Abcam); anti-GLUT1 (ab15309, Abcam); anti-c16orf62/VPS35L (PA5-28553, ThermoFisher Scientific); anti-SNX17(HPA043867, Atlas Antibodies); anti-SNX27 (ab77799, Abcam); anti CCDC93 (20861-1AP, Proteintech); anti-α-Tubulin (T9026, Sigma); anti-Vps26 (ab181352, Abcam); anti-VPS26C/DSCR3 (ABN87, Merck Millipore), WIPI1 (W2394, Sigma); WIPI2 (HPA019852, Sigma).

As secondary antibody we used: Cy3-conjugated Affinity Pure Donkey anti-Mouse IgG H + L (Jackson Immuno Research); Cy3-conjugated AffiniPure Donkey anti-Rabbit IgG H + L (Jackson Immuno Research); Alexa Fluor®488-conjugated AffiniPure Donkey anti-Rabbit IgG H + L (Jackson Immuno Research); Alexa Fluor®488-conjugated AffiniPure Donkey anti-Mouse IgG H + L (Jackson Immuno Research); Alexa Fluor® 647-conjugated AffiniPure donkey anti-rabbit IgG H + L (Jackson Immuno Research); Alexa Fluor® 647-conjugated AffiniPure donkey anti-Mouse IgG H + L (Jackson Immuno Research). Antibodies were used at dilutions summarized in Supplementary Table1.

Other reagents: Opti-MEM (Thermo Fischer, 11058021) and Trypsin (Thermo Fischer, 27250018).

Protease inhibitor (PI) cocktail final concentrations in samples: 100 μM pefablock SC (Merck, 11429876001), 2 μM leupeptin (Merck, 11529048001), 50 μM o-phenanthroline (Merck ,131377), 0.7 μM pepstatin A (Merck, 11524488001) dissolved in methanol.

Phosphatase inhibitor cocktail final concentrations: 10 mM NaF (S-7920, Sigma); 1mM Na Orthovanadate (S6508, Sigma); 2 mM β-glycerophosphate (A2253, AppliChem) all dissolved in water.

### Complementary DNA (cDNA) constructs

Vectors expressing tagged RAB proteins were purchased from Addgene: RFP-RAB4 (79,800; deposited by J.D. Johnson). DsRed-RAB11 (12679; deposited by R. Pagano); mCherry-RAB7 (61804; deposited by G. Voeltz); mCherry-RAB5 (49201; deposited by G. Voeltz). The following vectors were kindly provided by colleagues: EGFP-WIPI2 (pAR31CD vector) (Tassula Proikas-Cezanne, Tübingen, Germany). The sequences of WIPI1 and WIPI2 that have been used are given in Supplementary File 1.

### Site-directed mutagenesis

All constructs were verified by DNA sequencing. ^EGFP^WIPI2 or WIPI2^HA^ was used as DNA template for site-directed mutagenesis (QuikChange mutagenesis system, Agilent Technologies, 200524) to generate point mutations in the FSSS motif (S67A and S67E) following the manufacturer’s protocol.

To get WIPI2^S67E^ we used the following primers: Fw: AGATTGTTCTCCGAGAGCCTAGTGGCC and Rv GGC CAC TAG GCT CTC GGA

GAA CAA TCT; while for WIPI2^S67A^ we used Fw: AGATTGTTCTCCGCTAGCCTAGTGGCC

Rv: GGCCACTAGGCTAGCGGAGAACAATCT (Microsynth).

WIPI2^SLoop^ was generated by four consecutive amplifications using the following primers:

1) Fw: TGGACC GGGTACAAAGGGAAAGTG CTC

Rv: GAGCACTTTCCCTTTGTACCCGGTCCA;

2) Fw: GGGTACAAAGGGTTCGTGCTCATGGCC

Rv: GGCCATGAGCACGAACCCTTTGTACCC;

3) Fw: CTC ATG GCC TCC TAC AGC TAC CTG CCT

Rv: AGGCAGGTAGCTGTAGGAGGCCATGAG;

4) Fw: GCC TCC TAC AGC ACC CTG CCT TCC CAA
Rv: TTGGGAAGGCAGGGTGCTGTAGGAGGC

Non-mutated template vector was removed from the PCR mixture through digestion by the enzyme Dpn1 for 1 h at 37°C. The product was purified using NucleoSpin PCR Clean-up (Macherey-Nagel, 740609.50S) and transformed into *Escherichia coli*. Plasmid DNA was purified and sequenced. The site-directed mutagenesis system was also used to generate the siRNA-resistant cDNA for WIPI2 with primers Fw: CGATAGTCCTTTAGCCGCA and Rv: TGCGGCTAAAGGACTATCG. All oligonucleotides were synthesized by Microsynth.

### Cell culture, transfection and treatments

HK2 cells were grown in DMEM-HAM’s F12 (Thermo Fischer, 11765054) supplemented with 5% fetal calf serum (Gibco, 10270106), 50 U/mL penicillin/50 mg/mL streptomycin (ThermoFischer, 15140148), ITS (5 μg/mL insulin, 5 μg/mL TF, 5 ng/mL selenium; LuBio Science, 00-101-100ML). Cells were grown at 37°C in 5% CO_2_ and at 98% humidity.

HK2 cells were transfected with plasmids using the X-tremeGENE^TM^ HP DNA transfection reagent (Sigma-Aldrich, 6366546001), unless otherwise specified, according to the manufacturer’s instructions, and incubated for 18-24 h before fixation or live cell imaging. The HK2 cell line was checked for mycoplasma contamination by a PCR-based method. All cell-based experiments were independently repeated at least three times.

### Generation of HK2 cells expressing WIPI2^HA^

HK2 cell lines stably expressing the three forms (WT, S67A, S67E) of C-terminal HA-tagged WIPI2 were generated by lentiviral transduction. WIPI2 was cloned into modified pLKO.1 lentiviral vector (available from Sigma) between Xba1 and EcoR1 restriction sites. To obtain WIPI2^HA^ a sequence of 3Gly - 1Ser - 3Gly - 1Ser - HA - HA (designed and purchased from GeneScript) was added using Gibson assembly according to the manufacturer’s protocol. Finally, the lentiviral plasmid DNA was purified and sequenced. Lentiviral transfer vector together with third generation envelope and packaging plasmids were transfected into 293T cells to generate lentivirus. Packaged lentivirus containing supernatant was added to recipient HK2 cells in the presence of 10 μg/ml Polybrene (AL-118, Sigma-Aldrich). HK2 cells were stably infected with a lentiviral vector for constitutive WIPI2^HA^ expression for 6 h. 2 days after infection, cells were selected with 1 μg/ml of puromycin for 3 days. Protein samples were collected and used for IP experiments 7 days after infection.

### RNA interference

HK2 cells were transfected with siRNA for 72 h using Lipofectamine® RNAiMax (Thermo Fisher Scientific, 13778150) according to the manufacturer’s instructions. Control cells were treated with identical concentrations of siGENOME Control Pool Non-Targeting from Dharmacon (D-001206-13-05). siRNAs targeting WIPI2 were from Dharmacon (ON-TARGETplus Human WIPI2 J-020521). siRNAs were used at a final concentration of 20 nM.

### Endocytosis assays

#### EGFR degradation

HK2 cells were seeded in DMEM-HAM’s F12 supplemented with 5% foetal calf serum, 50 U/mL penicillin/50 mg/mL streptomycin and ITS (5 μg/mL insulin, 5 μg/mL TF, 5 ng/mL selenium. The day after the growth medium was replaced with the same but without serum for 24 h and HK2 cells were stimulated with 100 ng/ml EGF for the indicated periods of time. The cells were fixed for 10 min in 4% paraformaldehyde in PBS at different time points after stimulation for IF studies (Figure 1 A-B) or washed and lysed for immunoblotting analysis (Figure D)

### Immunofluorescence

Cells were fixed for 10 min in 4% paraformaldehyde in PBS. After fixation, cells were permeabilized in 0.05% (w:v) saponin (Sigma-Aldrich, 558255), 0.5% (w:v) BSA and 50 mM NH_4_Cl in PBS (blocking buffer) for 30 min at room temperature. The cells were incubated for 1 h with primary antibodies in blocking buffer, washed three times in PBS, incubated for 1 h with the secondary antibodies (Alexa Fluor-conjugated), washed three times in PBS, mounted with fluorescence mounting medium (Dako, S3023) on slides and analysed by confocal microscopy.

### Confocal fluorescence microscopy, image processing, and colocalization analysis

HK2 cells were grown to 70% confluence on glass coverslips and immunofluorescence microscopy was performed as described above. The experiments were repeated at least three times and representative images are shown. Colocalization was analysed by acquiring serial sections from about 100 cells per sample. Images were exported in TIFF format and processed as previously described (Vicinanza et al., 2011). Images of samples to be compared were acquired using the same settings (i.e. laser power, photomultiplier gain and pinhole size), avoiding pixel saturation. The images were processed in the same way using ImageJ software. Channels from each image were converted into 8-bit format and the ‘Auto Local Threshold’ Plug-in with the ‘Default’ method was used to segment grayscale images, identify the structures of interest and subtract background.

GLUT1, β1-Integrin, EEA1, and LAMP1 fluorescence levels were quantified using ImageJ. *z*-stack images were acquired and compressed into a single plane using the ‘maximum intensity Z-projection’ function in Image J. Individual cells were selected using the freeform drawing tool to create a ROI (ROI). The ‘Measure’ function provided the area, the mean grey value and integrated intensity of the ROI. The mean background level was obtained by measuring the intensity in three different regions outside the cells, dividing them by the area of the regions measured, and averaging the values obtained. The background was subtracted from each cell to obtain the CTCF (corrected total cell fluorescence). It was calculated using the formula: CTCF=integrated intensity of cell ROI − (area of ROI × mean fluorescence of background).

To quantify the degree of colocalization, confocal z-stacks were acquired, but a single panel was used for the measurement. Single channels from each image in 8-bit format were thresholded to subtract background and then the "Just Another Colocalisation Plug-in" (JACoP) ImageJ software was used to measure the overlap coefficient according to Manders. Manders’ coefficient indicates the overlap of the signals and represents the degree of colocalization:

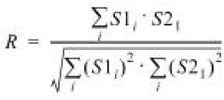

where *S*1 represents signal intensity of pixels in the channel 1 and *S*2 represents signal intensity of pixels in the channel 2. The colocalization coefficients M1 and M2 indicate the contribution of each channel to the pixels. They are independent of the intensity of the overlapping pixels, but they are sensitive to background, so a threshold must be set (Manders et al., 2011).

For the triple colocalization analysis (Figures S9-S11) we first performed colocalization between two single channels (i.e., red and blue) and then took that subset of overlapping spots (either a binary image or a list of spots) to measure how many colocalized with green signals.

To measure colocalization of RAB-proteins on tubules in cells overexpressing ^EGFP^WIPI2 variants (Figure S4) we used an approach based on the detection of signal edges through the “Canny or Sobel filters” Plug-in for ImageJ (Canny, 1986) . The surrounded fields were filled to generate binary mask images. In this way, we were able to select tubules in order to separate signal from background, but also to determine a common region for analyzing both channels (i.e., red for Rabs and green for WIPI2). These mask images were then subjected to thresholding as described above and colocalization was evaluated with the ImageJ JACOP Plug-in by calculating Manders’ coefficient. Confocal microscopy was performed on an inverted confocal laser microscope (Zeiss LSM 880 with airyscan) with a 63x 1.4 NA oil immersion lens unless stated otherwise.

### Gel electrophoresis and Western blot

Control cells were plated into 12-well tissue culture test plates (TPP) until they reached around 80-90% confluency, except for knockdown-cells, which were cultured until 72h after transfection with the siRNAs by reaching a confluence of 80%. Cells were then washed three times with ice-cold PBS (phosphate-buffered saline), scraped, and proteins were extracted in ice-cold lysis buffer (150 mM NaCl, 2 mM EDTA, 40 mM HEPES-NaOH pH 7.4, and 1% Triton X-100 supplemented with phosphatases and protease inhibitor cocktail (see details in Reagent section). Protein extracts were supplemented with 1/4 volume of 5x reducing sample buffer (250 mM Tris-Cl, pH 6.8, 5% β-mercaptoethanol, 10% SDS, 30% glycerol, 0.02% bromophenol blue) and heated to 95 °C for 5 min. The samples were run on either 8%, 10% or 12.5% SDS-polyacrylamide gels (W x L x H: 8.6 x 6.8 x 0.15 cm). The stacking gels were prepared as follows: 6% acrylamide, 0.16% bis-acrylamide, 0.1 M Tris, pH 6.8, 0.1% SDS, 0.1% TEMED, 0.05% ammonium persulfate. Running gels were: 10% or 12.5% acrylamide, 0.27% or 0.34% bis-acrylamide, 0.38 M Tris, pH 8.8, 0.1% SDS (Applichem, 475904-M), 0.06% TEMED (Applichem, A1148), 0.06% APS (Applichem, A2941). The gels were run at constant current (20–30 mA). Proteins were blotted onto nitrocellulose membrane by the semi-dry method for 80 min at 400 mA (Trans-Blot® SD Semi-Dry Electrophoretic Transfer Cell, Bio-Rad). After incubation with the primary antibody, signals were detected by secondary antibodies coupled to infrared dyes (LI-COR) and detected on a LI-COR Odyssey infrared fluorescence imager. Images were exported as TIFF files and processed in Adobe Photoshop. Band intensity was quantified using ImageJ band analysis.

### Immunoprecipitation of Retriever

HK2 cells expressing the three variants (WT, S67E, S67A) of WIPI2 tagged with HA were grown to 80% confluence in 15 cm dishes. After rinsing the cells three times with phosphate buffered saline (PBS 1X), they were detached and lysed by incubating 15 min at 4°C with lysis buffer: 150 mM NaCl, 2 mM EDTA, 40 mM HEPES-NaOH pH 7.4, 1% Triton X-100, supplemented with phosphatases and proteases inhibitor cocktails (details in Reagent section) and 1 mM PMSF out of a 200 mM solution in ethanol, which was freshly prepared immediately before use. The lysate was spun for 10 min at 10,000 x g on a cold bench-top centrifuge). 2.5% of the clarified extracts was reserved as “input”. Each clarified extract was incubated for 2 h at 4°C with 20 μl of Anti-HA High Affinity antibody (clone 3F10) (Roche #118150116001). Then, samples were washed three times with lysis buffer and, after discarding the last wash, the beads were resuspended in 50 μl of pre-warmed elution buffer (2% SDS, 1mM EDTA, 10mM DTT, 20mM Tris HCl pH 7.4, 100 µM PMSF) and shaken at 55°C for 10 min. Samples were centrifuged, the supernatants were transferred to new Eppendorf tubes, and 5x sample buffer containing 100 mM of fresh DTT was added. Samples were kept at 95°C for 5 min and centrifuged in a table to centrifuge. All supernatants were transferred to a new Eppendorf tube yielding the “IP” samples. Inputs and IP were loaded on 8% or 10% SDS–polyacrylamide gels.

### Statistical analysis

Differences between the means have been evaluated by an unpaired Student’s t-test assuming unequal variances unless otherwise specified. They are considered as significant differences when p < 0.01. Experiments were independently repeated at least three times. For fluorescence microscopy quantifications at least 30 cells were analysed from each experiment. Representative images are shown. Western blotting experiments were independently repeated at least three times and representative blots are shown. Means ± standard deviation (s.d.) are shown unless specified otherwise.

## Data and reagent availability

Primary data, clonal reagents and cell lines will be supplied by the authors upon request.

## Supporting information

Supplementary table of reagents

## Acknowledgements

We thank Navin Gopaldass and Thibault Courtellemont for discussions. This work was supported by grants from the SNSF (31003A_179306, 310030_204713, and 10.006.083) and ERC (788442) to AM.

---

**Figure. S1.**
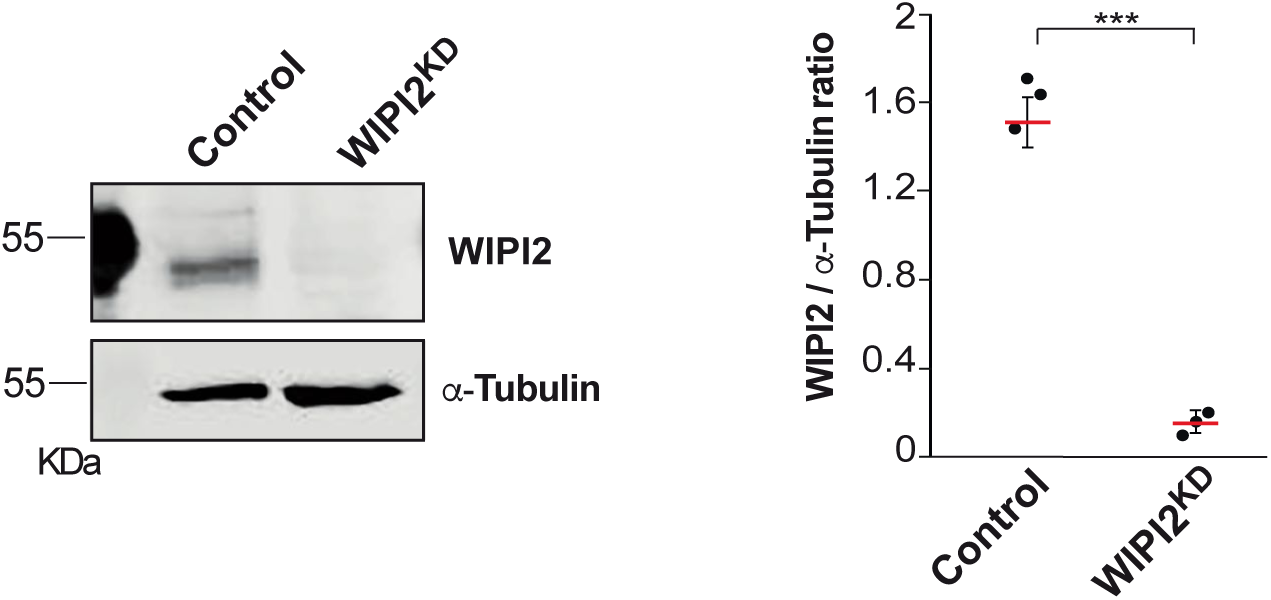
Efficiency of WIPI2 knockdown. Lysates of HK2 cells (50 μg per sample) were treated with non-target siRNA (Control) or siRNA against WIPI2 (WIPI2^KD^). HK2 cells were analyzed by SDS-PAGE and Western blot against the indicated proteins. α-tubulin was used as a loading control. A representative blot is shown. The signals were quantified on a LICOR Odyssey infrared fluorescence image WIPI2/a-Tubulin ratios were calculated. Red bars indicate the mean and error bars the SEM; n=3 independent biological experiments. P-values were calculated by an unpaired Student’s t-test with unequal variances. ***P < 0.0001.

**Figure S2.**
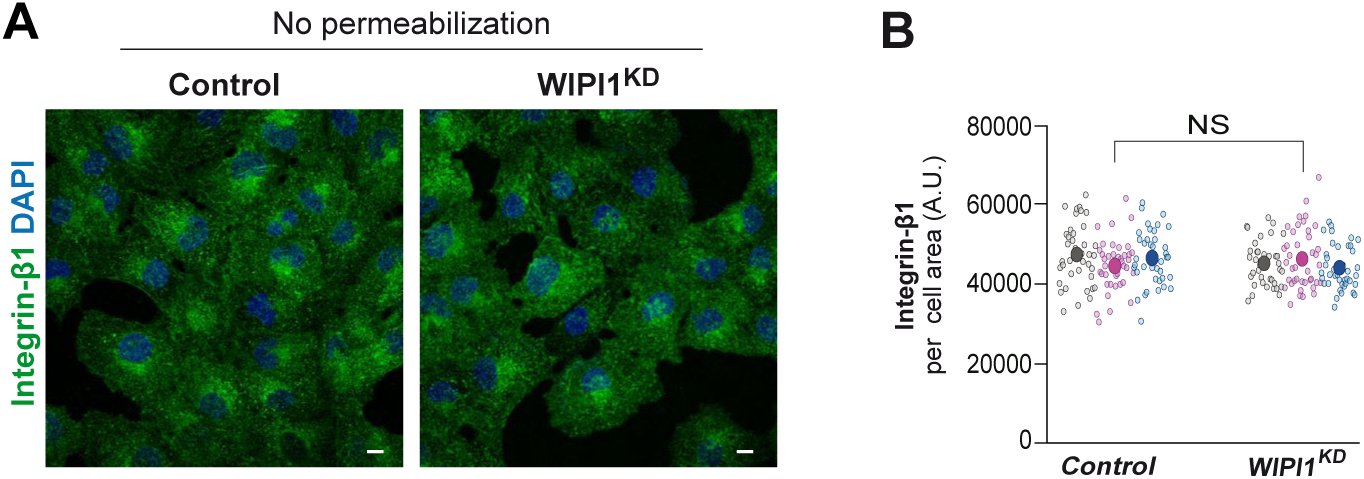
Surface expression of β1-Integrin in cells depleted of WIPI1. **A.** Control and WIPI1^KD^ cells were fixed and stained with DAPI and with antibody to β1-Integrin. Scale bars: 10 μm. **B.** Quantification of β1-Integrin-immunofluorescence in cells from A. Regions of interest (ROIs) corresponding to each cell and in some regions outside the cells (background) were manually defined using ImageJ software. Total cell fluorescence was integrated, and the background fluorescence was subtracted. The resulting total cell fluorescence was divided by the area of the cell. This value is shown in the graph. 150 cells per condition, stemming from three independent biological experiments, were quantified. Individual cells and means are presented by smaller and larger circles, respectively, coloured according to the independent experiment that they stem from. P values were calculated applying an unpaired Student’s t-test with unequal variances The analysis was performed with 99% confidence. NS = not significant (P>0.05).

**Figure S3.**
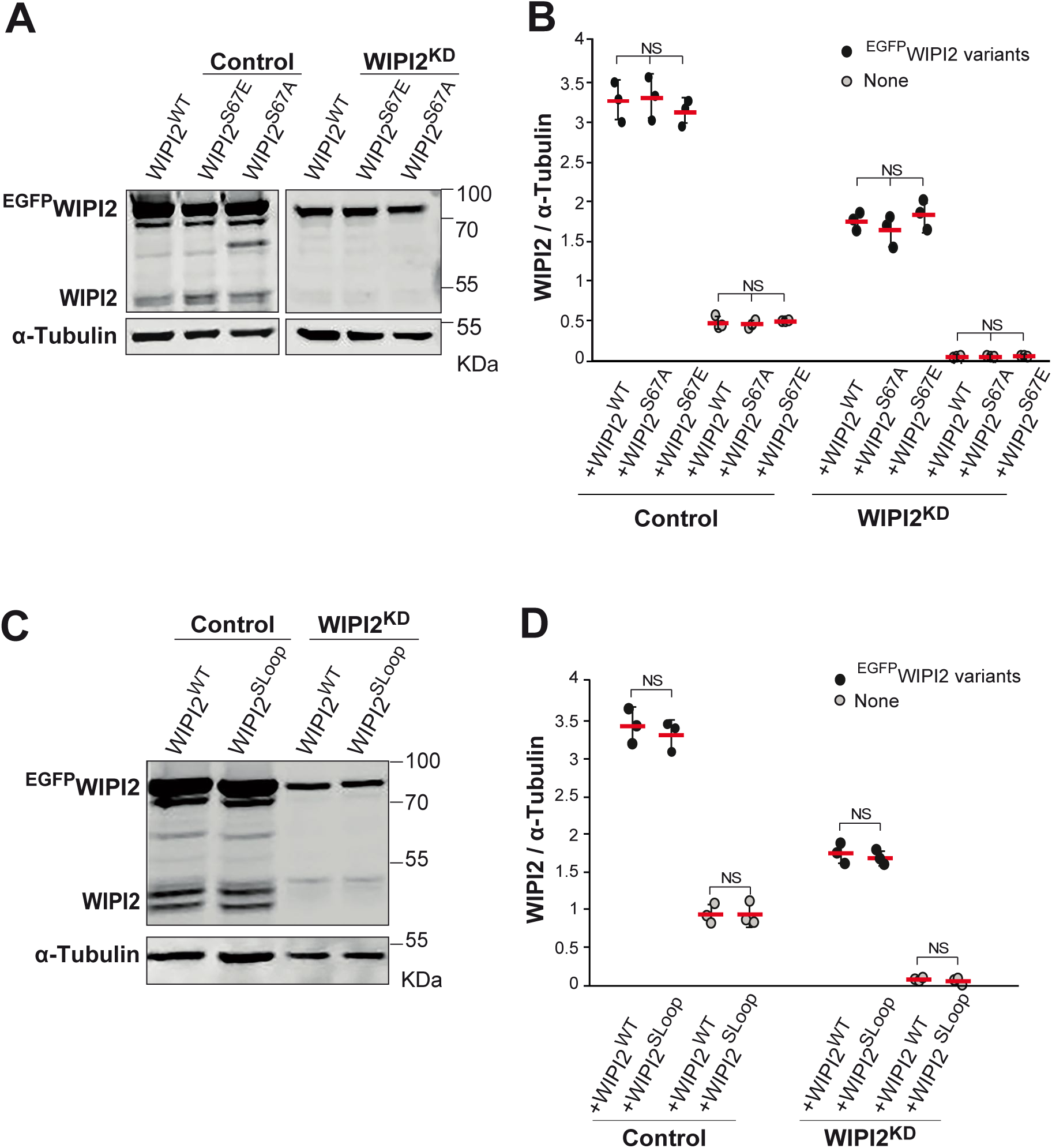
Expression levels of WIPI2 variants. **A.** FSSS variants. Cell lysates from Control and WIPI2^KD^ cells were transfected for 18 h with siRNA resistant constructs for ^EGFP^WIPI2^WT^, ^EGFP^WIPI2^S67A^ and ^EGFP^WIPI2^S67E^. The cells were analyzed by SDS–PAGE and Western blotting with antibodies to WIPI2 and α-Tubulin. A representative blot is shown. **B.** Quantification of the signals for transiently expressed ^EGFP^WIPI2 variants from A and for endogenous WIPI2. Blots from three independent biological experiments were quantified on a Licor Odyssey infrared scanner. Graphs on the right side show the mean and SEM. P-values were calculated using an unpaired Student’s t-test with unequal variances. NS: not significant. **C.** SLoop variant. ^EGFP^WIPI2 and ^EGFP^WIPI2^SLoop^ were expressed and analyzed by Western blotting as in described in A**. D.** Blots from C were quantified as in B.

**Figure S4.**
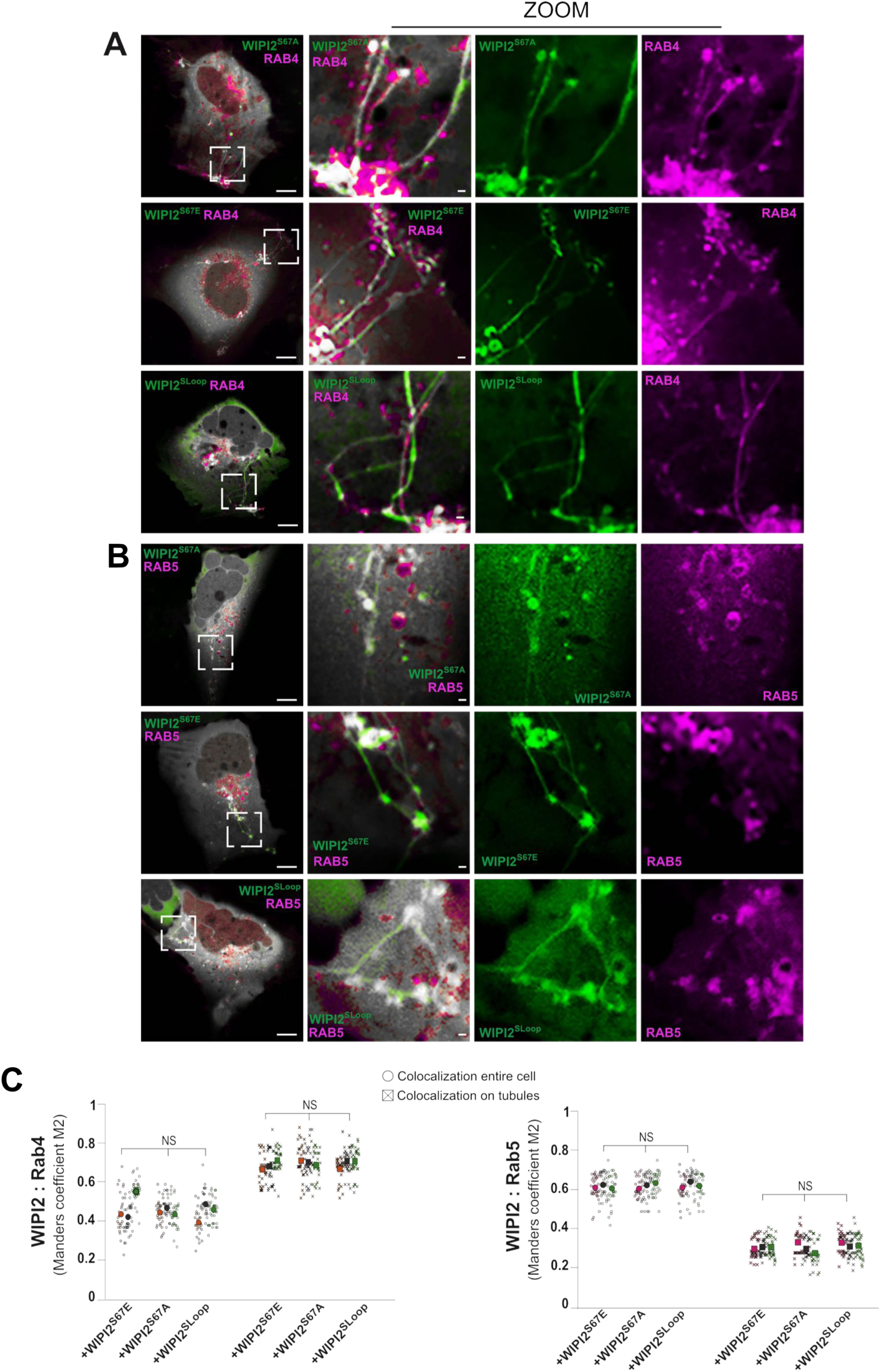
^EGFP^WIPI2 variants are preferentially associated with recycling endosomes. **A.B.** Colocalization with RAB proteins. HK2 cells were transiently transfected with plasmids carrying the indicated ^EGFP^WIPI2 variants with mCherry-RAB5 or RFP-RAB4. After 18h transfection, cells were analysed by live cell confocal microscopy. Arrows point to tubular structures that are readily detectable in cells expressing the SLoop and S67 variants of WIPI2. Scale bars: 10 μm. Insets show enlargements of the outlined areas with a white dashed line. Scale bars: 2 μm. **C.** Colocalization analysis of the images from (A-B). Quantification was carried out for the tubular structures and the entire cell using the Manders’ colocalization coefficient M2, indicating the fraction of green pixels overlapping with magenta pixels, calculated in Image J software. Colocalization was quantified from three independent experiments with a total of 90 cells per condition. Values of individual cells and means are presented by smaller and larger circles, respectively, coloured according to the independent experiment that they stem from. An unpaired Student’s t-test with unequal variances was used to calculate P-values. The analysis was performed with 99% confidence. NS = not significant (P>0.05).

**Figure S5.**
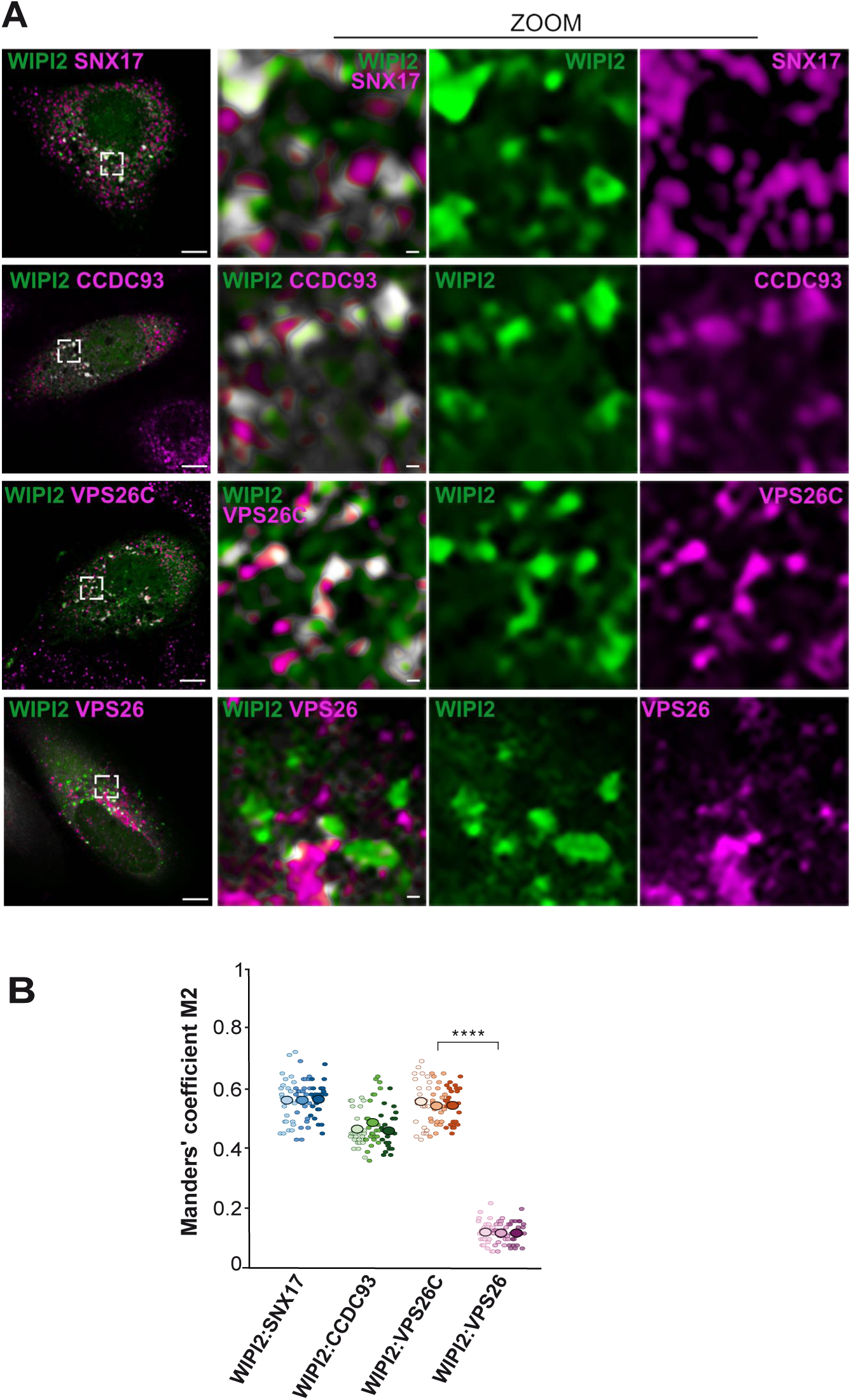
WIPI2 colocalization with Retriever, CCC complex and Retromer. **A.** WIPI2 colocalization with SNX17, CCDC93, VPS26C and VPS26. HK2 cells were fixed 18 h after transient transfection with ^EGFP^WIPI2, stained with the indicated antibodies and imaged by confocal microscopy. Scale bars: 10 μm. Insets show enlargements of the outlined areas with a white dashed line. Scale bars: 2 μm. **B.** Quantification. Using the images from (A), colocalization was assessed using Manders’ colocalization coefficient M2, calculating the fraction of green pixels overlapping with magenta pixels. 75 cells per condition, stemming from three independent biological experiments, were quantified. Values of individual cells and means are presented by smaller and larger circles, respectively, coloured according to the independent experiment that generated them. P values were calculated applying an unpaired Student’s t-test with unequal variances. The analysis was performed with 99% confidence. ****P < 0.00001.

**Figure S6.**
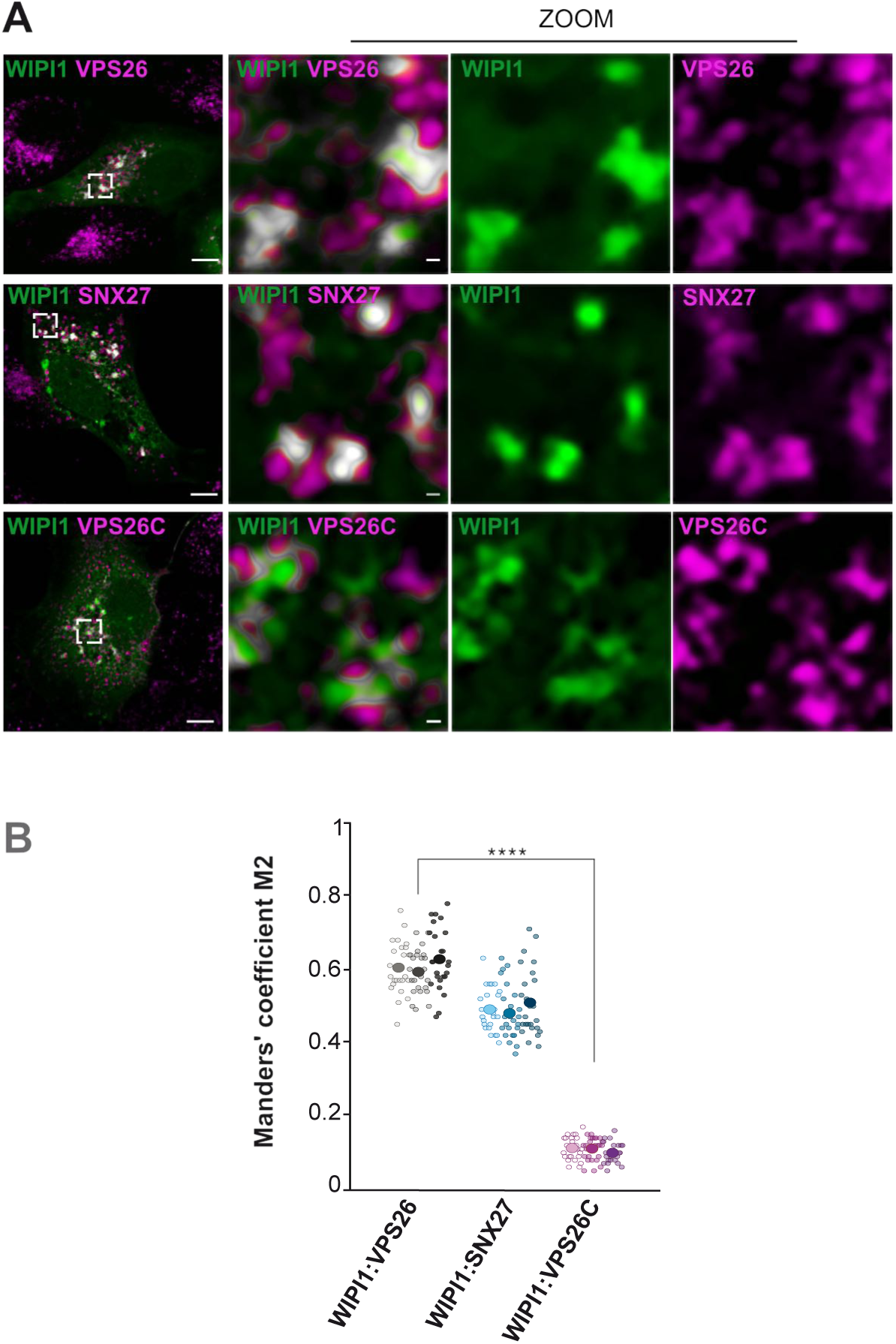
WIPI1 colocalizes with VPS26, but not with VPS26C. **A.** WIPI1 colocalization with VPS26, SNX27, and VPS26C. HK2 cells were fixed 18 h after transient transfection with ^EGFP^WIPI1, stained with the indicated antibodies and imaged by confocal microscopy. Scale bars: 10 μm. Insets show enlargements of the outlined areas with a white dashed line. Scale bars: 2 μm. **B.** Quantification. Using the images from (A), colocalization was assessed using Manders’ colocalization coefficient M2, indicating the fraction of green pixels overlapping with magenta pixels, calculated in Image J software. 75 cells per condition, stemming from three independent biological experiments, were quantified. Individual cells and means are presented by smaller and larger circles, respectively, coloured according to the independent experiment. P values were calculated applying an unpaired Student’s t-test with unequal variances. The analysis was performed with 99% confidence. ****P < 0.00001.

**Figure S7.**
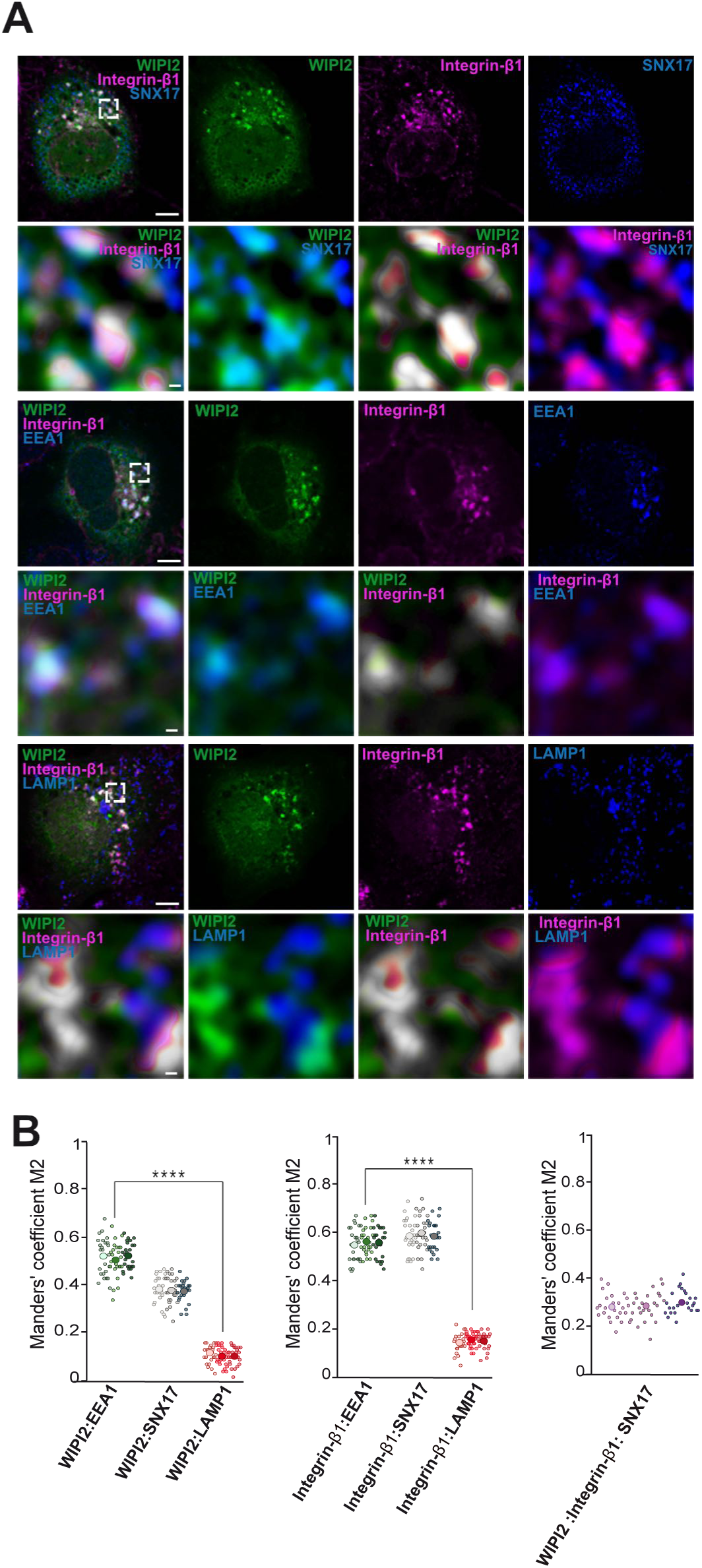
WIPI2 colocalizes with Integrin-β1 on early endosomal compartments. **A.** WIPI2 colocalization with Integrin-β1. HK2 cells were fixed 18 h after transient transfection with ^EGFP^WIPI2, stained with the indicated antibodies and imaged by confocal microscopy. Scale bars: 10 μm. Insets show enlargements of the outlined areas with a white dashed line. Scale bars for these are 2 μm. **B.** Quantification. Using the images from (A), colocalization was assessed using Manders’ colocalization coefficient M2. 75 cells per condition, stemming from three independent biological experiments, were quantified. Individual cells and means are presented by smaller and larger circles, respectively, coloured according to the independent experiment. P values were calculated applying an unpaired Student’s t-test with unequal variances. The analysis was performed with 99% confidence. ****P < 0.00001.

**Figure S8.**
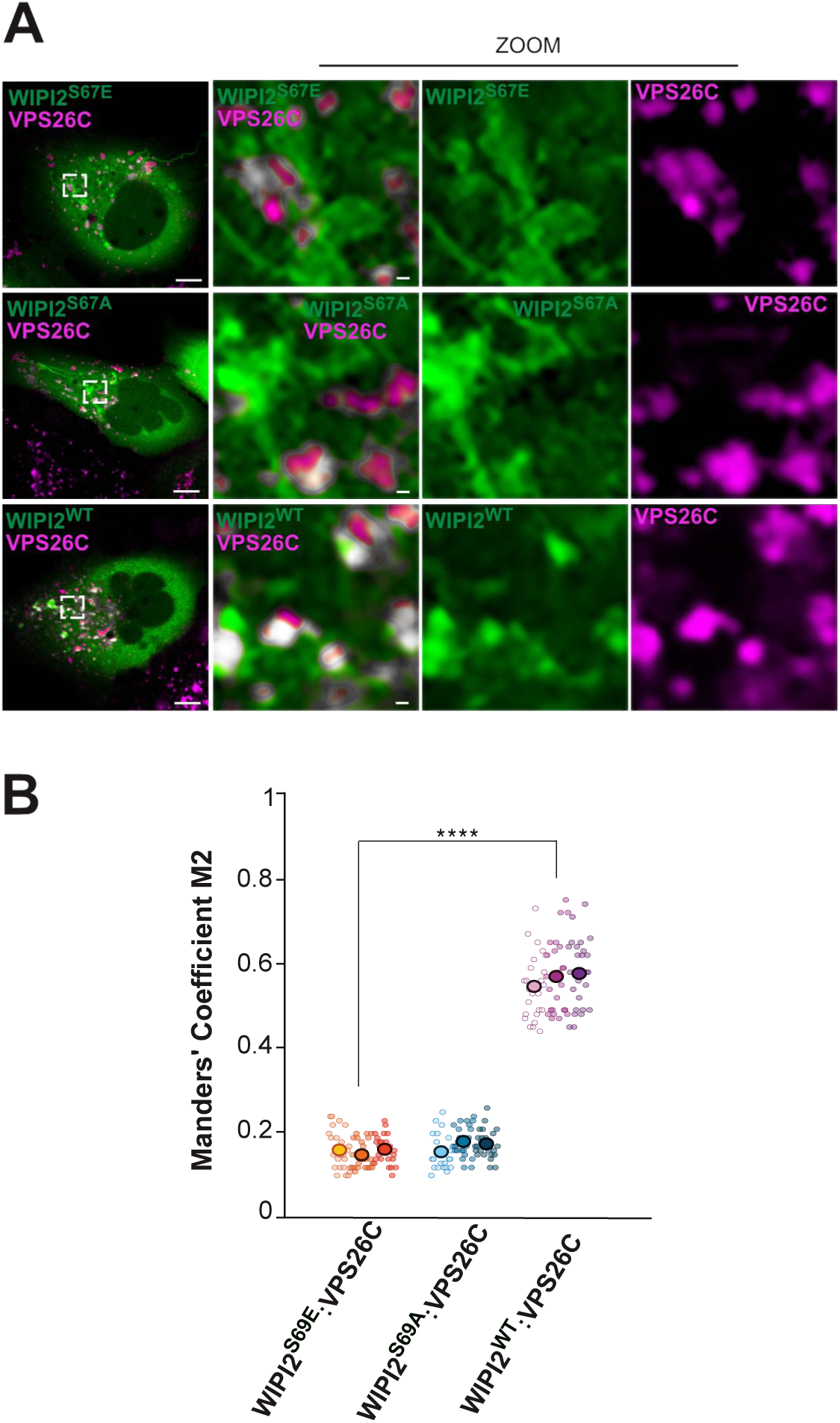
Impact of the FSSS motif on colocalization of WIPI2 and Retriever. **A.** HK2 cells were fixed 18 h after transient transfection with the indicated ^EGFP^WIPI2 variants, stained with antibodies to VPS26C and imaged by confocal microscopy. Scale bars: 10 μm. Insets show enlargements of the outlined areas with a white dashed line. Scale bars: 2 μm **B.** Quantitative colocalization analysis of the images from (A). Quantification was assessed using the Manders’ colocalization coefficient M2, indicating the fraction of green pixels overlapping with magenta pixels, calculated in Image J software. 75 cells per condition, stemming from three independent biological experiments, were quantified. Values of individual cells and means are presented by smaller and larger circles, respectively, coloured according to the independent experiment that generated them. P values were calculated applying an unpaired Student’s t-test with unequal variances. The analysis was performed with 99% confidence. ****P < 0.00001.

**Figure S9.**
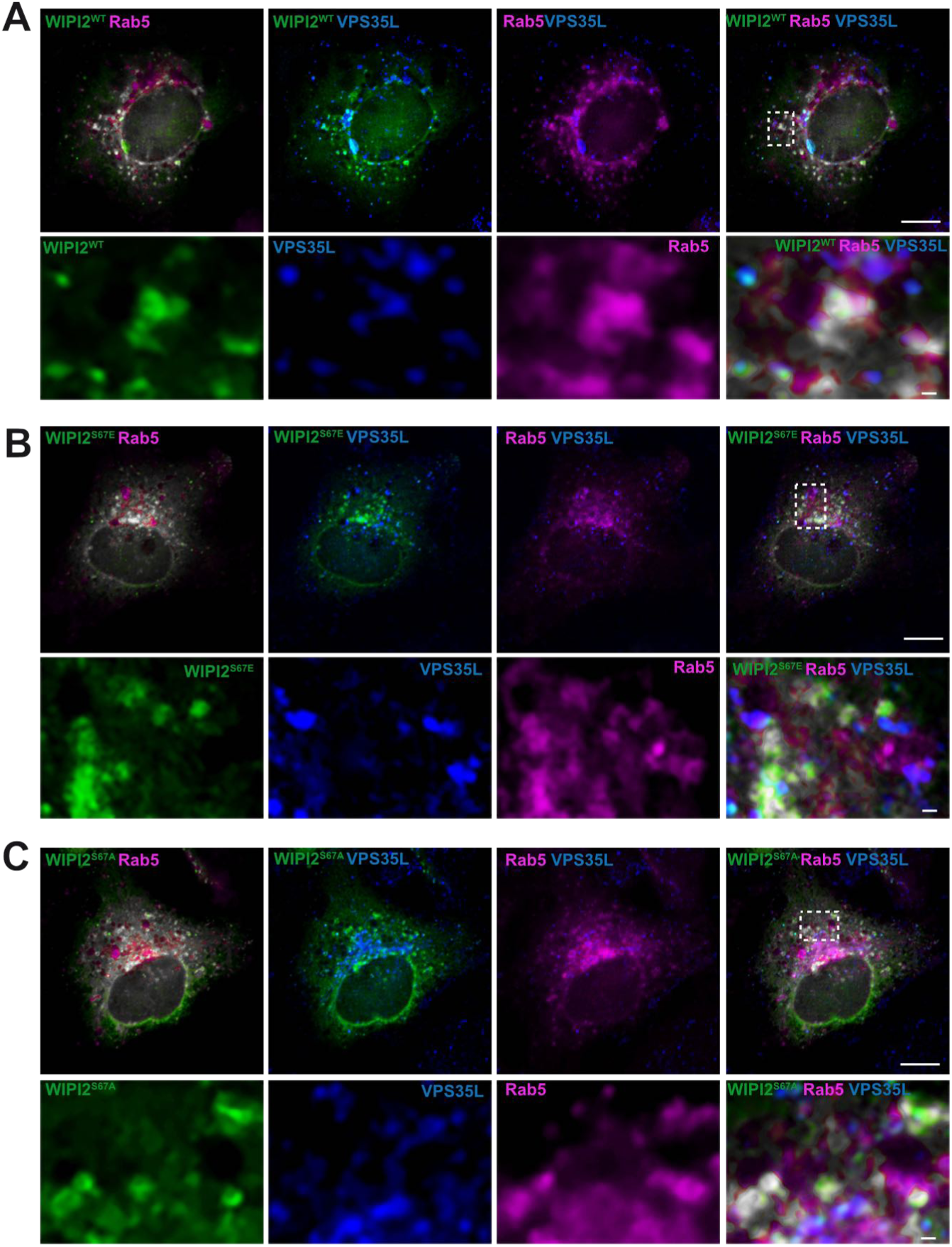
WIPI2 colocalization with RAB5 and Retriever. HK2 cells expressing mCherry-RAB5 and ^EGFP^WIPI2 wild-type (**A**) or its indicated variants (**B.C**), were fixed, permeabilized and stained with antibodies to VPS35L (blue). Scale bars: 10 μm. Insets show enlargements of the outlined areas. Scale bars: 2 μm. Quantifications from multiple experiments are shown in Figure. 9

**Figure S10.**
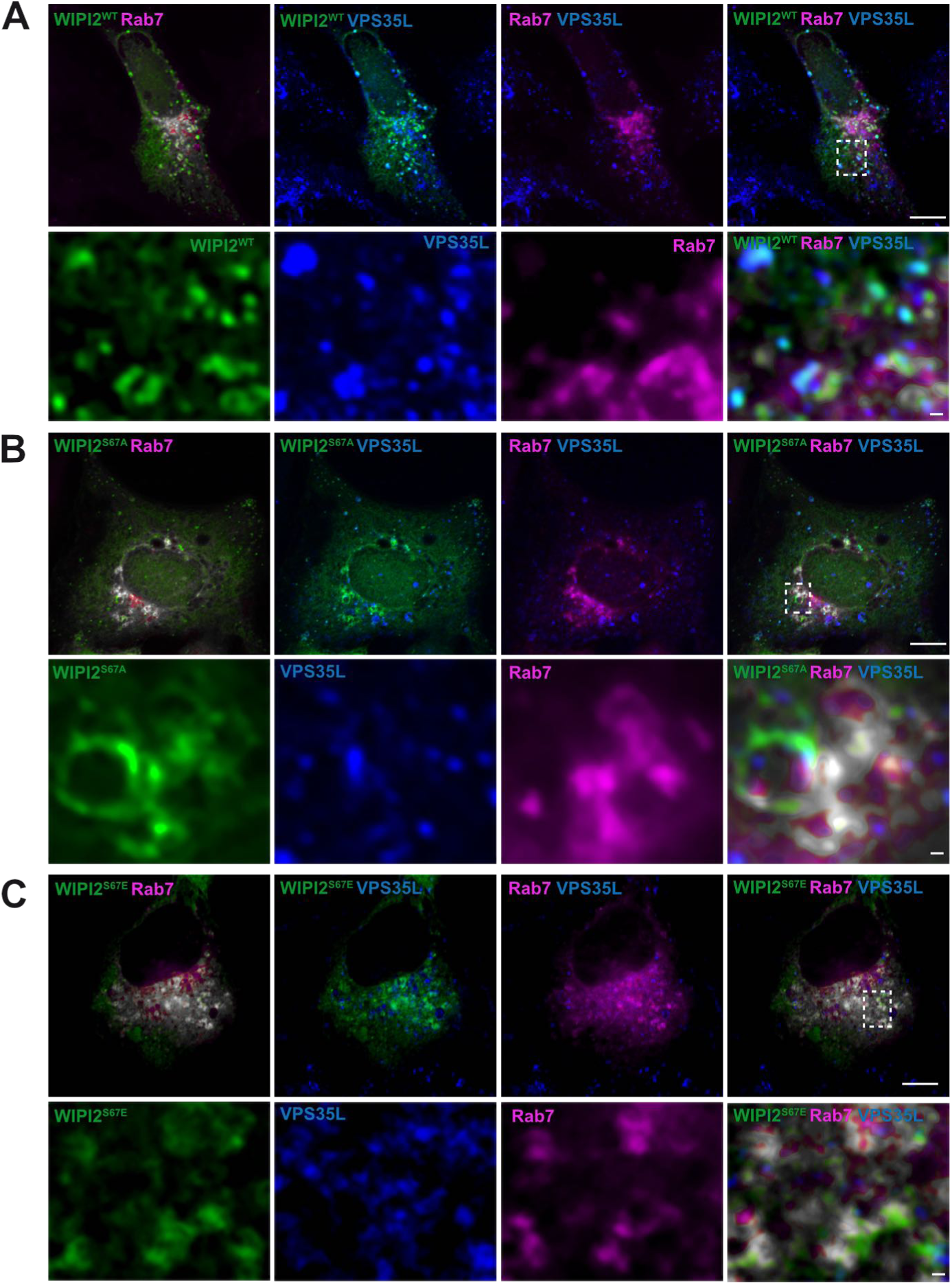
WIPI2 colocalization with RAB7 and Retriever. HK2 cells expressing mCherry-RAB7 and wild-type ^EGFP^WIPI2 (**A**) or its indicated variants (**B.C**) were fixed, permeabilized and stained with antibodies to VPS35L (blue). Scale bars: 10 μm. Insets show enlargements of the outlined areas. Scale bars: 2 μm. Quantifications from multiple experiments are shown in Figure. 9

**Figure S11.**
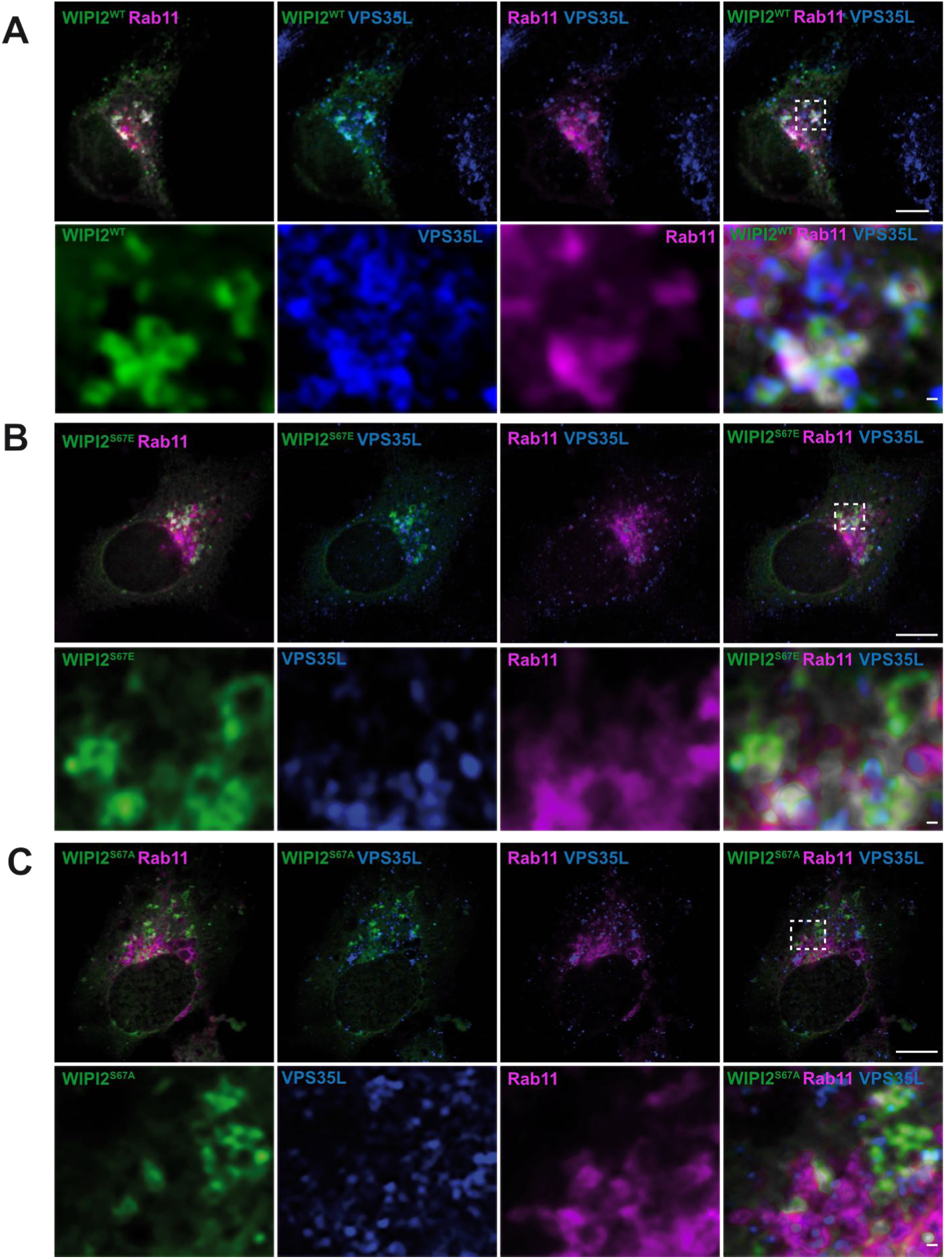
WIPI2 colocalization with RAB11 and Retriever. HK2 cells expressing mCherry-RAB7 and wild-type ^EGFP^WIPI2 (**A**) or its indicated variants (**B.C**) were fixed, permeabilized and stained with antibodies to VPS35L (blue). Scale bars: 10 μm. Insets show enlargements of the outlined areas. Scale bars: 2 μm. Quantifications from multiple experiments are shown in Figure. 9

## Supplementary File 1

Sequences of WIPI1 and WIPI2 that had been used for mutagenesis and for generating tagged alleles.

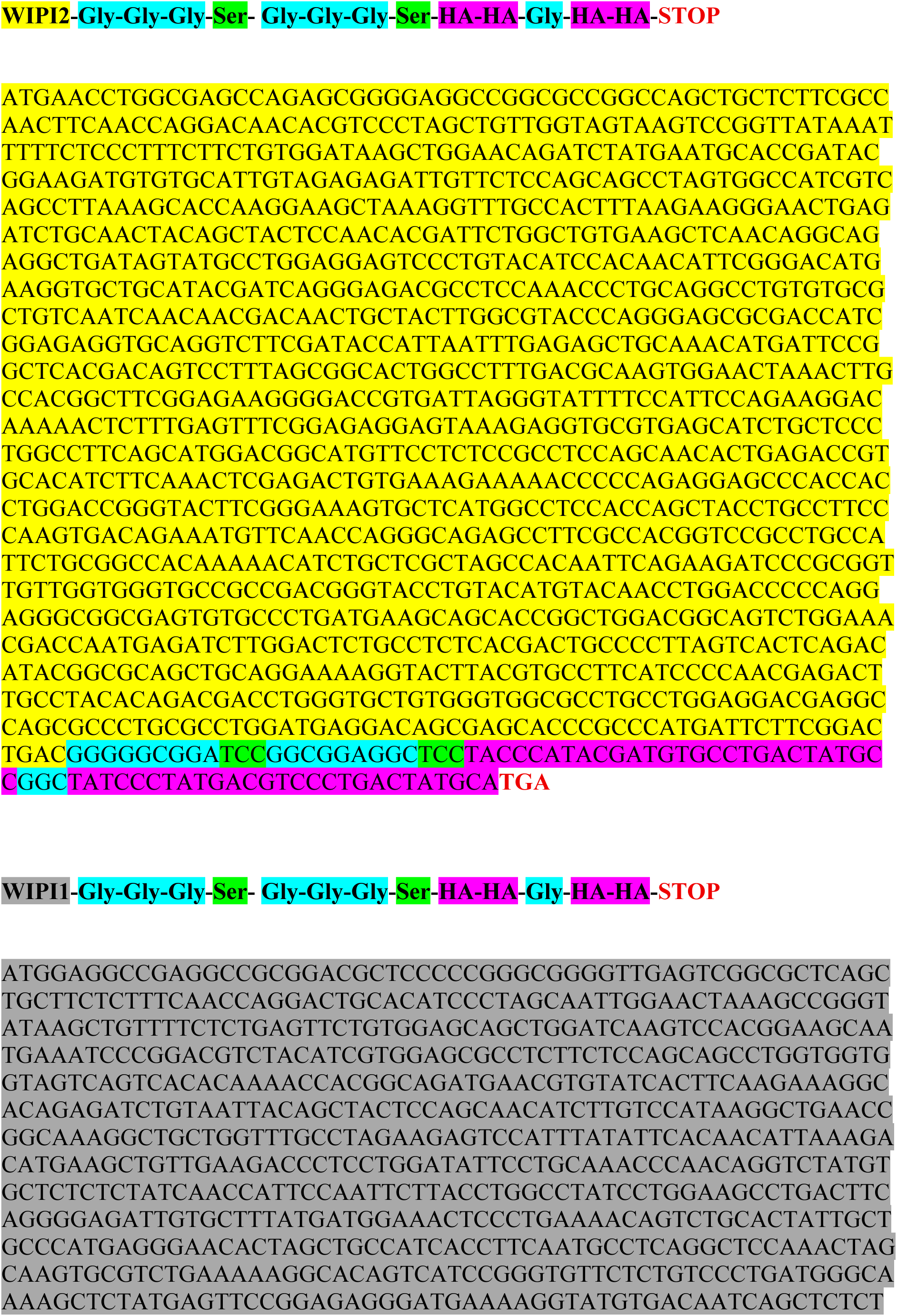
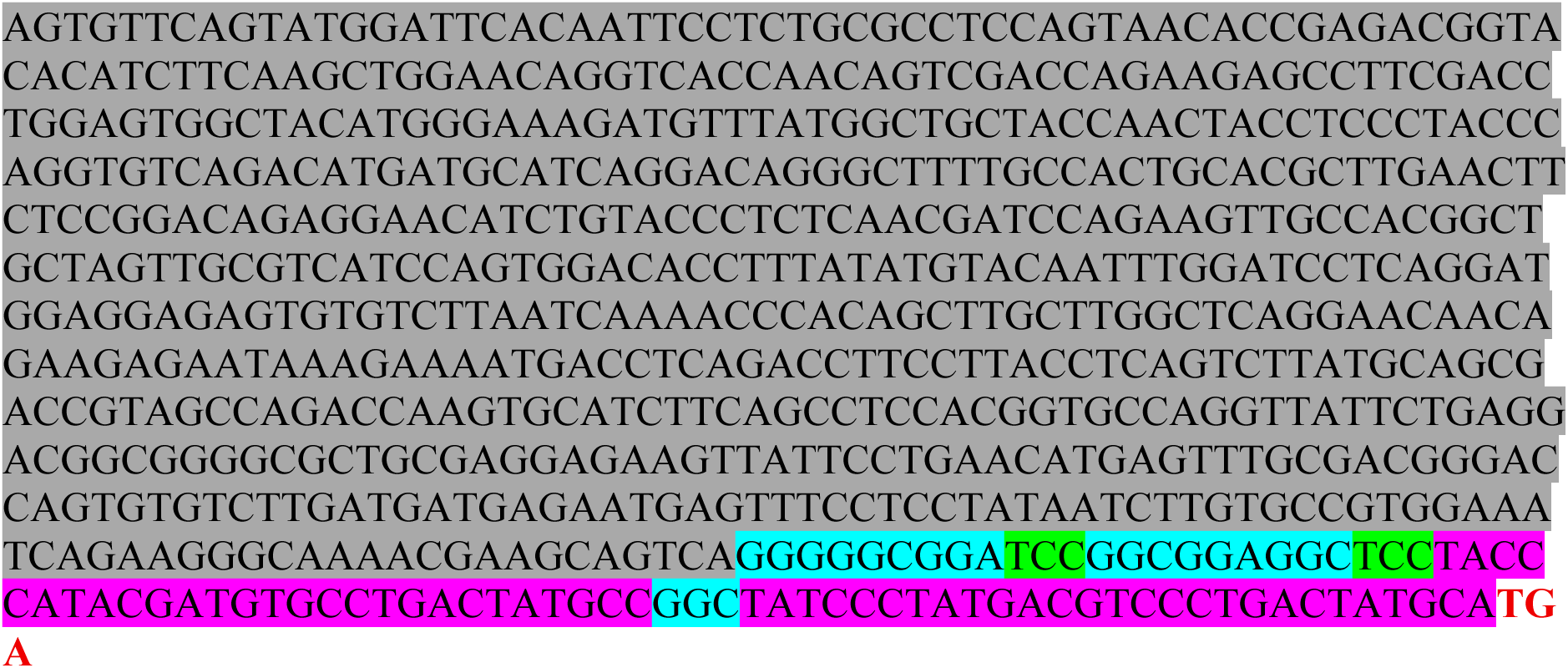

## Notes

### Competing Interest Statement

The authors have declared no competing interest.

### Summary of Updates

Improved represenation of individual data points; better description of statistical analysis; better images; additional data.

